# Nutrient sensing; transcriptomic response and regulation of gut motility in an agastric vertebrate

**DOI:** 10.1101/827659

**Authors:** Hoang T. M. D. Le, Kai K. Lie, Angela Etayo, Ivar Rønnestad, Øystein Sæle

## Abstract

The transcriptome of nutrient sensing and the regulation of gut motility by nutrients in a stomachless fish with a short digestive tract; the ballan wrasse (*Labrus berggylta*) were investigated. Using an in vitro model, we differentiate how signals initiated by physical stretch and nutrients modulate the gut evacuation rate and motility patterns, and transcriptomic changes. Stretch on the intestine by inert cellulose initiated fast evacuation out of the anterior intestine compared to the digestible protein and lipid. Stretch on the intestine upregulated genes associated with increased muscle activity, whereas nutrients stimulated pathways related to ribosomal activity and the increase in the expression of several neuropeptides which are directly involved in gut motility regulation. Our findings show that physical pressure in the intestine initiate contractions propelling the matter towards the exit, whereas the sensing of nutrients modulates the motility to prolong the residence of digesta in the digestive tract for optimal digestion.

**Summary statement:** Pressure by food speed up peristalsis in the intestine, but the intestines ability to sense nutrients slow down peristalsis for better digestion. This is partly controlled by genetic regulation.

## 1. Introduction

When food is ingested, senses such as vision, smell and taste initiate a cascade of signals that prepare the body, and the digestive system in particular, for the tasks ahead. Digestion and absorption of nutrients are facilitated via a combination of mechanical force (mixing and transport) produced by gut motility and chemical breakdown by digestive enzymes. The mixing and propulsion of gut contents along the alimentary cannal and the evacuation of unabsorbed particles are performed by strong smooth muscular contractions observed as various motility patterns (Chang and Leung, 2014). Motility patterns are classified into two main categories: *i*) non-propulsive contractions, such as segmentation, and *ii*) propulsive contractions, such as peristalsis and migrating motor complexes (MMCs) (Holmgren and Olsson, 2009; Welcome, 2018). Each type of contraction plays a specific role, depending on its amplitude, propagation distance and velocity. Standing contractions (Brijs et al., 2014; Brijs et al., 2017) also called stationary contractions (Huge et al., 1995) or segmentations (Gwynne and Bornstein, 2007b; Gwynne et al., 2004; Hennig et al., 2010) are a set of stationary and rhythmic contractions of the circular muscle, which was first described by Cannon more than 100 years ago (Cannon, 1902; Cannon, 1912). This contraction type participates in the digestion (e.g. mixing ingesta with digestive juice) and absorption (i.e. exposing digested gut contents to the absorptive epithelium) (Andrews and Young, 1993; Cannon, 1912; Chang and Leung, 2014; Janssen et al., 2011). Propulsive contractions, also referred to as peristalsis or propagating contractions, are a series of annular contractions that produce force to propel luminal contents distally (Cannon, 1912; Chang and Leung, 2014; Ehrlein et al., 1983; Ehrlein et al., 1987; Szurszewski, 1969). The propagating contractions are classified into two main sub-types based on their amplitude and velocity. Ripples are defined as a set of rhythmic and shallow contractions, which propagate for a relatively short distance and at high speed in either the anterograde or retrograde direction (Brijs et al., 2014; D’Antona et al., 2001). It has been suggested that this type of contraction mixes the luminal content for digestion and absorption rather than propelling it (D’Antona et al., 2001). Another sub-type of propulsive contraction is characterized as a series of high-amplitude contractions that propagate for a longer distance than ripples but at a low velocity. In some fish studies these are called slow propagating contractions (Brijs et al., 2014; Brijs et al., 2017; Brijs et al., 2016; Le et al., 2019a), while in studies of humans and other mammals, the accepted term is “high-amplitude propagated contractions” (Bharucha, 2012; Spiller et al., 1988). These move luminal contents from one section to the next, thanks to their high amplitude (Bharucha, 2012; Husebye, 1999; Kunze and Furness, 1999).

Since different types of contraction serve different functions, the motility patterns may change as a function of the caloric level or composition of the digesta passing through the gastrointestinal tract, and also on feeding status. Post-prandial segmentation is stimulated and modulated by nutrients present in the gut and the intestinal transit rate is slowed down. In contrast, the presence of inert / non-nutritive ingesta in the intestinal lumen abolishes segmentation and stimulates propulsive contractions; thus, the intestinal evacuation rate increases to egest the indigested content (Borgstrom and Arborelius, 1975; Defilippi and Gómez, 1995; Ehrlein et al., 1987; Keinke and Ehrlein, 1983; Schmid and Ehrlein, 1993; Siegle and Ehrlein, 1988). Motility patterns differ between feeding and fasting periods. In mammals, when the stomach and intestine are emptied after a meal, the motility shifts to typical myoelectrical patterns that consist of three or four phases, namely the MMCs. This contraction type is made up of a sequence of contractile waves that clean and propel digestive waste, unabsorbed particles and microbiota in the anal direction (Deloose et al., 2012; Deloose and Tack, 2015; Sarna, 1986; Szurszewski, 1969). In fish, high frequency of standing contractions with or without ripples, was observed in fed shorthorn sculpin, whereas when the fish was fasting, slow propagating contractions and retrograde ripples frequently occurred (Brijs et al., 2014).

As in other vertebrates, the gastrointestinal mucosa in fish encompass a large number of enteroendocrine cells (EEC) and sensory systems. Two components of the sensory system are mechanoreceptors (sensory receptors that respond to mechanical pressure or distortion) and chemoreceptors (receptors that transduce a chemical substance and generate biological signals) (Bertrand, 2009; Lowe and Anderson, 2015). The gastrointestinal hormones/neuropeptides are involved in both local signaling pathways like the enteric nervous systems within the alimentary canal and remote signaling pathways that originate in the central nervous system. The enteric nervous system mainly regulates gut motility via afferent, efferent and interneurons that connect smooth muscle layers of the gut wall and the parasympathetic and sympathetic nervous systems (Kunze and Furness, 1999; Olsson and Holmgren, 2001).

The gut responses to nutrients, both physically (i.e. changes in gut motility induced by food) and at molecular level (i.e. changes in expression of genes related to digestion, absorption and metabolism), as have been well described in mammals (e.g. Defilippi and Gómez, 1995; Ehrlein et al., 1987; Gwynne and Bornstein, 2007a; Hurst et al., 2014; Miguel-Aliaga, 2012; Schwartz, 2011; Siegle and Ehrlein, 1988; van der Wielen et al., 2014). The impact of nutrient sensing on food intake regulation and energy metabolism has been widely investigated in fish (e.g. Babaei et al., 2017; Comesana et al., 2018; Comesaña et al., 2018; Jiang et al., 2017; Li et al., 2017; Tian et al., 2018; Xu et al., 2016), and has been reviewed by Conde-Sieira and Soengas (2017). However, effects of nutritive and non-nutritive meals on gut motility and intestinal epithelium metabolism in fish remains largely unknown, except for a study in motility patterns in fed and starved sculpin (*Myoxocephalus scorpius*) (Brijs et al., 2014). The recent study aimed to determine how different nutrients regulate both gut motility and gene expression in the intestine. We analyzed changes in motility patterns of isolated whole gut preparations administered a bolus of either a nutritional composition (viscous protein or lipid) or a bolus of an inert suspension containing cellulose or plastic beads. We used ballan wrasse juveniles as a model. This fish does not have a stomach and the whole intestine was used for the study. Differences in gene expression were compared between intestines administered cellulose versus empty intestines in order to isolate the effect of mechanical sensing; and intestines administered with either protein or lipid versus cellulose to compare effects of chemical sensing on enterocyte metabolism. We also described postprandial motility patterns as responses to the administration of the compounds in the ballan wrasse intestines.

## 2. Materials and Methods

### 2.1. Nutrient bolus

Six different diets, including intact lipid (IL), hydrolyzed lipid (HL), intact protein (IP), hydrolyzed protein (HP), cellulose (CL) and plastic beads (PB) were prepared. Intact lipid was made by stirring a mix of 80% (by volume) cod liver oil (Møllers Tran, containing omega-3-fatty acids and Vitamin D), 15% phosphate-buffered saline (PBS) pH = 8, and 5% Tween 20 (TWEEN® 20, P9416 Sigma Aldrich) to obtain the same viscosity as the hydrolyzed lipid diet. To make 5 mL hydrolyzed lipid diet, we incubated 4 mL cod liver oil with 3.5 mg lipase (Lipase from *Pseudomonas cepacia* - 62309 Sigma Aldrich, ≥30 U/mg) dissolved in 750 µL PBS (pH = 8) at 40°C for 5 hours. The pH of the mix was maintained at around 8 by titrating with 5 M NaOH during incubation. The mix was then incubated at 80°C for 2 hours to deactivate the lipase before mixing with 250 µL Tween. 20. Five mL of intact protein diet was prepared by combining 2 g casein with 4 mL 100 mM NH_4_HCO_3_ and 1 mL 1 mM HCl. Hydrolyzed protein diet was prepared by incubating 5 mL intact protein diet with 43.7 mg trypsin (Trypsin Powder, Porcine 1:250, 85450C, Sigma Aldrich) at 37°C for 20 hours, followed by 80°C for 2 hours for enzyme deactivation. 5 mL of hydrolyzed protein diet was freshly mixed with 100 µL protease inhibitor [Protease/Phosphatase Inhibitor Cocktail (100X) #5872, Cell Signaling Technology, Inc, The Netherlands] before administration of the bolus into the intestine. Cellulose diet was created using 0.5 mg cellulose (Cellulose microcrystalline powder, 435236 Sigma Aldrich) in 1.5 mL H_2_O, adding a tiny amount of Brillant Blue (Brilliant Blue R, B7920 Sigma-Aldrich) as a color maker. These five bolus types were made and stored at – 20°C for use within a week. The diets were thawed and warmed up at 14°C before being administered into the intestines.

### 2.2. Animals and tissue preparation

Ballan wrasse juveniles were supplied by a commercial fish farm (Marine Harvest Labrus, øygarden, near Bergen, Norway). The fish were reared in accordance with the Norwegian Animal Welfare Act of 12 December 1974, no. 73, §§22 and 30, amended 19 June 2009. The facility has general permission to rear all developmental stages of *Labrus berggylta*, license number H øN0038 provided by the Norwegian Directorate of Fisheries (https://www.fiskeridir.no/English). Fish were nursed in 3 m^3^-tanks in a temperature-controlled room (around 14°C) under a 24:0 h light:dark photoperiod and fed every 15 minutes with a commercial pellet diet. Fish weighing 20 - 30 g were transferred from the Marine Harvest farm to the Institute of Marine Research (Bergen, Norway) laboratory and were kept under conditions identical to those at the nursing station for one day prior to the experiments.

On the day of the trial, the fish were anesthetized in 0.05 mg/mL tricaine methanesulfonate (MS222) dissolved in sea water prior to euthanasia spinal cord lesion using a scalpel and removal of the intestine. The eviscerated intestine included esophagus and anus with the surrounding skin, leaving the whole intestine intact. Eviscerated intestines (5 – 9 cm length) were immediately immersed in Ringer’s solution according to Rønnestad et al. (2000) with some modifications (in mM: NaCl, 129; KCl, 2.5; MgCl_2_, 0.47; CaCl_2_, 1.5; NaHCO_3_, 20.2; and NaH_2_PO_4_, 0.42). The intestines were administered a bolus of one of the five diets (IL, HL, IP, HP, and CL were described in section *Nutrient bolus*) or one plastic bead (2 – 3 mm diameter and 20 – 30 mg weight) to mimic ingestion of a meal of 0.1% body weight (Le et al., 2019b). The prepared intestines were rapidly mounted in individual glass tubes containing 25 mL of Ringer’s solution at 14°C, aerated with 95% O_2_ and 5% CO_2_. Thereafter, the intestines were carefully stretched out longitudinally inside the tube with the oral opening closed and the anus open according to Le et al. (2019a) and incubated for 14 hours or 1 hour depending on the protocol for each of the two experiments (Figure 1) below.

**Figure 1.**
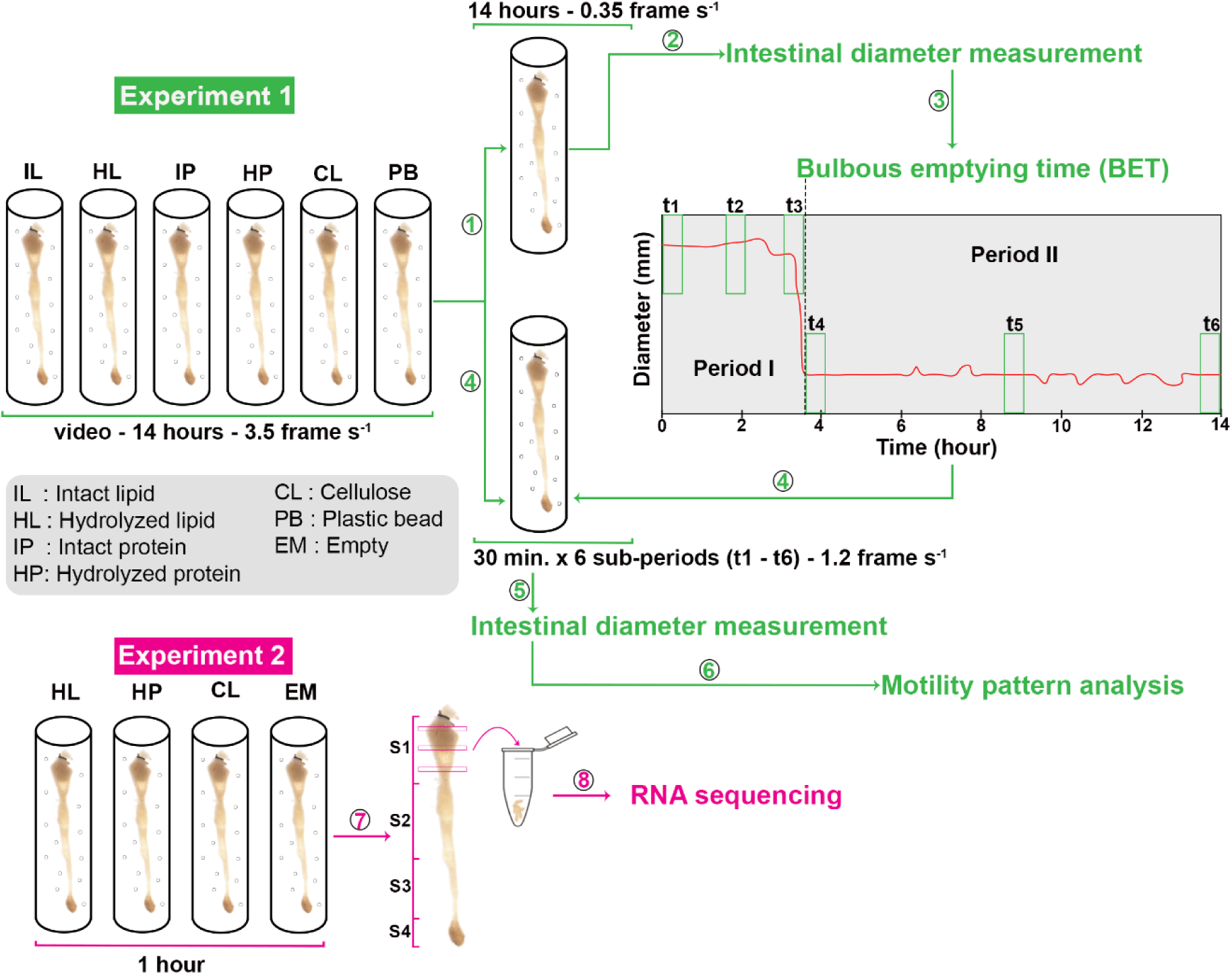
Experimental design and data analysis. Experiment 1 analyzed the effects of nutrient bolus on gut motility. Each intestine was administered a bolus of one of the five diets (IL, HL, IP, HP, and CL) or one plastic bead (PB) before mounting in an individual glass tube containing Ringer’s solution with air supply. A time-lapse series of images of the intestines was captured at intervals of 3.5 frame s^-1^ for 14.5 h. ➀, every tenth frame was subtracted from the original videos (3.5 frames s^-1^) to obtain 0.35 frame s^-1^ videos and ➁, intestinal diameters along the intestine on each image frame were measured using NIS-Elements Confocal 4.51.01 software. ➂, the data of intestinal diameter was used to calculate the time of emptying of the bulbous (BET). Based on BET, times for six sub-periods were defined to select the frames to be in step ➃. A recovery period of 20 min. was set after mounting the intestines to the glass tube. The period analyzed was from 0 h on the horizontal axis of the Diameter and Time graph, which was set at the 21^st^ min. after the recovery period until 14 h. The sub-period t1 was the first 30 min. after the recovery period; t2 for 30 min. between the 0 h and the BET; t3 for the last 30 min. before the BET; t4, t5 and t6 were the first, middle, and last 30 min., respectively, of the period after the BET until 14 hours after the 0h point. ➃, every third frame was subtracted from the original videos for the six sub-periods. ➄, intestinal diameter was measured on the video frames for each sub-period (t1 – t6) and this data was used to analyze motility patterns in ➅. Experiment 2 was to collect gut tissue samples for gene expression analysis. Three nutrient bolus types (HL, HP, CL) were selected for this experiment. The isolated intestines were administrated a bolus of three selected diets and incubated in Ringer’s solution for 1 h before tissue samples of the bulbous were collected ➆ and analyzed for RNA sequencing ➇.

### 2.3. Experiment 1: Description of motility patterns

#### 2.3.1. Image acquisition

Based on our observations, movements within the intestines occurred 10 – 20 min. after they had been mounted in the medium. Thus, the activity of the intestines was recorded for a total of 14 hours after a 20-min.-acclimatation period to examine the effects of the different nutrients on intestinal motility. The 14-hour duration was chosen for Experiment 1 according to the *in vivo* passage rate of ballan wrasse juveniles, in which ingested feed took 10 – 14 h to pass through the alimentary tract (Le et al., 2019b). A time-lapse series of images of intestines was captured during the experiment using a camera (Nikon DS-Fi3) with a macro lens (Nikon, AF Micro-Nikkor 60mm f/2.8D), at a resolution of 1024×768 pixels. The capture of the time-lapse series was controlled with the NIS-Elements Confocal 4.51.01 software and captured 3.5 frames s^-1^. In this experiment, we used six dietary treatments (IL, HL, IP, HP, CL, and PB) with six replicates, and six intestines were processed in parallel in each video. The videos were then used to determine the time at which the bolus was transferred from the bulbous - Segment 1 to the next intestinal sections and to examine the motility patterns of the intestines (Experiment 1, Figure 1)

#### 2.3.2 Time of emptying the bulbous (Segment 1)

The bulbous is a morphologically distinct feature of the anterior gut of the ballan wrasse, with an expanded diameter relative to more posterior regions [Figure 1 in Le et al. (2019b)]. Here we defined the time of emptying the bulbous (bulbous emptying time - BET) as the time at which Segment 1 propelled the chyme to the next section and reduced its average diameter to its minimum value. First, we measured the diameters along the intestines during 14 h. The method used to quantify changes in width along the whole length of the intestines over time has been fully described in Le et al. (2019a). Briefly, every tenth frame was extracted from the original videos (3.5 frames s^-1^) to obtain 0.35 frame s^-1^ videos (➀ in Figure 1) and calibrated for length and time before being analyzed using the NIS-Elements Confocal 4.51.01 software. A threshold for intensity was manually selected to cover the whole intestinal area on each frame. Background noise, generated by air bubbles and equipment accessories, was removed using the “restrictions” functions in the Nis-Element software, whereby the size of the intestine is defined on the basis of pixel recognition and the program removes items in images outside of the intestine. The diameter of the intestine was measured at every one pixel using the function “Automated measurement” in the NIS-Elements software and a diameter-matrix was produced. The diameter-matrix, which is a numerical matrix of diameter along the intestine for 14 hours was used to generate spatio-temporal (ST) maps and to calculate the average width of Segment 1. In this study, we defined the four ballan wrasse intestine segments as the average ratios of segment length / total intestine length for Segment 1 (bulbous), Segment 2, Segment 3, and Segment 4 (hindgut), i.e. 0.39, 0.23, 0.23, and 0.15, respectively, according to the morphological description by Le et al. (2019b) (on Figure 1) and Le et al. (2019a). The average width of Segment 1 (AW1) in each frame in the 14-hour video was calculated as the mean value of diameters at the pixels covering a length of 39% of the anterior intestine. The 5^th^ percentile value of the AW1 for 14 hours (α) was defined using “quantile” function in R [quantile (data of AW1 in 14 h, 0.05)]. The empty bulbous was assumed to represent an average width which was equal to or smaller than α and named as “bulbous emptying width – BEW”. The BEW values occurred at various points in time during the 14-hour period. Thus, the time for emptying the bulbous (BET) was defined as the time when the first BEW value occurred after starting to record the treatments.

BET was first examined in three fish using intestinal diameters which were measured at five interval values of 3.5, 1.2, 0.7, 0.35, and 0.02 frames s^-1^ for 14 hours. The results for BET evaluated in the videos at equal or more than 0.35 frames s^-1^ did not differ. The ST maps constructed from videos at 0.35 frames s^-1^ represented well the pattern of change in intestinal width. Thus, we measured intestinal diameter in 14-hour videos at 0.35 frames s^-1^ in order to examine BET. All BETs were verified on the time-lapse videos and the ST maps.

#### 2.3.3. Analysis of motility patterns

Gut motility patterns were analyzed for two defined periods: period I was from 0 h (i.e. 20 min. after the insert of a bolus) to BET, and period II was from BET to 14h post-starting point. Motility patterns in each period were analyzed from three 30 min. sub-periods based on the experimental design in (Brijs et al., 2014; Brijs et al., 2017). The first sub-period (t1, Figure 1) was the first 30 min. of the recording. The second was the 30 min. between 0 h and the BET (t2, Figure 1). The third covered the last 30 min. before the BET (t3, Figure 1). Contractions defined within the three sub-periods t1 – t3 were used to analyze motility patterns (frequency, amplitude, duration, propagation direction, distance, and velocity) for period I, when ingesta remained at the bulbous (Segment 1). The fourth (t4), fifth (t5) and sixth (t6) sub-periods (Figure 1) were the first, middle, and last 30 min., respectively, of the period after the BET until 14 hours after the 0-h point. For the intestines that had a BET less than or equal to 1.5 hours, all frames within 0 h and BET were selected for the analysis of motility patterns.

Video frames for each intestine were extracted for six (or four for a BET of less than 1.5 hours) sub-periods and at intervals of 1.2 frames s^-1^ [according to the test by Le et al. (2019a)]. Intestinal diameters were measured along the intestines (as mentioned in the section *Time of emptying the bulbous (Segment 1)*, above) for analysis of motility patterns. In a recent study, we classified the motility patterns of ballan wrasse intestines into three types of contractions (standing contractions, ripples, and slow propagating contractions) according to Le et al. (2019a). We defined the types of contractions and their parameters (i.e. frequency, amplitude, propagating direction, distance, duration, and velocity) according to the description by Le et al. (2019a). Frequency of contractions (contractions per min at every mm length of intestine - cpm) and propagation direction (the proportion of contractions, propagating in either anterograde or retrograde direction, compared to the total number of contractions) in each period were calculated based on the total number of contractions that occurred within the three sub-periods in each intestinal segment. The remaining parameters (amplitude, distance, duration, and velocity) were presented as a median of the data set of all contractions during the three sub-periods.

#### 2.3.4. Data analysis

There were no differences in either motility patterns or emptying time between IP and or HL and IL. Furthermore, the analysis of free fatty acids and amino acids in the feces collected from intestines after 14 hours did not show a significant difference in nutrient composition between intact and hydrolyzed treatments (Figures S1 and 2). We therefore pooled the data from IP and HP into a group named “protein” and IL and HL into a group “lipid”; and omitted the intestines which looked deadly. BET and motility patterns were thus analyzed for four groups of nutrient boli (protein n = 12, lipid n = 11, cellulose n = 6, and plastic bead n = 6) for two periods (I and II) in four intestinal segments. One-way ANOVA followed by Tukey HSD were used to evaluate the effect of the four bolus treatments on time for BET. A linear mixed models (lme) analysis followed by Tukey HSD were used to compare the frequencies of each contraction type between the four treatment groups and the two defined periods. Amplitude, propagating distance and direction of each contraction type (continuous proportions ranging from 0 to 1) were treated as quasi-binomial response variables and compared, taking treatment or period as predictor variables for each intestinal segment using a generalized linear mixed models (glmmPQL) (R, version 3.4.2 released 2017-09-28, within R studio interphase (version 1.1.383) for Windows. Duration and velocity were treated as exponential variables in glmmPQL models. The lme and glmmPQL models included individual fish IDs as a random factor and the response variables (contraction parameters) as repeated measurements. Changing the contrast where each treatment/period became the intercept of the model was applied to determine the variation in parameter between the four treatments or between the two periods. Differences were treated as significant at p < 0.05 for all tests in this study. Amplitude, duration, propagation distance, and velocity are presented as median±s.d., and other data as mean±s.d.

### 2.4. Experiment 2: Transcriptome sequencing

The analysis of Experiment 1 showed that there was no difference in the transit time and motility between the intact (IL or IP) and pre-digested (HL or HP) meal. Nor were there differences in the composition of the free fatty acids or amino acids in the feces collected from intestines during the 14 hours between the intact and hydrolyzed treatments (Figures S1 & 2). We therefore selected HL, HP and CL diets to examine the transcriptomic effects of digestible and indigestible ingesta. We also analyzed gene expression in the empty intestines as a control measure. The HL and HP were employed to examine how the gut senses nutrients compared to the non-nutritive meal. The indigestible CL was used to evaluate the stretching effects on gut activity compared to the control and to separate the effects of stretching to that of nutrients in the HL and HP groups. The intestines fed 0.1% body weight, according to Le et al. (2019b) of one of three nutrient bolus types (HL, HP, or CL) and empty intestines were incubated in Ringer’s solution for one hour before tissue samples for transcriptomic analysis were collected. Segment 1 was selected for transcriptomic analysis because it is the main site for digestion (Le et al., 2019b). The first segment of the anterior part (about 40% of the total gut length) was cut off, opened by incision and gently washed with Ringer’s solution. The rest of the gut was discarded. Three pieces of tissue (around 20 mg each) were cut transversely from three parts of the first segment and placed in RNAlater^®^ (Sigma-Aldrich, Missouri, USA) for further RNA extraction (Experiment 2, Figure 1).

#### 2.4.1. RNA sequencing

Total RNA was extracted and treated with BioRobot® EZ1 and RNA Tissue Mini Kit (Qiagen, Hilden, Germany) as previously described by Le et al. (2019b). RNA quality and integrity were validated using a NanoDrop ND-1000 UV–vis Spectrophotometer (NanoDrop Technologies, Wilmington, USA) and Agilent 2100 Bioanalyzer and RNA 6000 Nano LabChip kit (Agilent Technologies, Palo Alto, USA) respectively. All samples had 260/230 and 260/280 ratios above 2.0 and 2.2 respectively. The average RNA integrity number (RIN) of all samples was 7.9±0.7. Sequencing and library preparation were performed by the Norwegian Sequencing Centre (www.sequencing.uio.no). DNA libraries were prepared as previously described by Le et al. (2019b) using 90 ng total RNA input to the TruSeq Stranded mRNA Library Prep Kit (Illumina, San Diego, California, USA). For multiplexing, standard Illumina adaptors were used. The libraries were sequenced using the NextSeq Illumina platform (Illumina, San Diego, California, USA) according to the manufacturer’s instructions, generating single end 75bp read libraries with an average library size of 25±6 million reads. Raw reads were submitted to the gene expression omnibus https://www.ncbi.nlm.nih.gov/geo/ (accession number GSE129459).

#### 2.4.2. Differential gene expression analysis

Adaptor removing and quality trimming was performed using the TrimGalore 0.4.2 wrapper tool and default parameters. Library quality was investigated using fastQC embedded in the TrimGalore wrapper (https://github.com/FelixKrueger/TrimGalore). Each intestinal RNAseq library was mapped individually to the labrus genome assembly (European Nucleotide Archive accession number: PRJEB13687, http://www.ebi.ac.uk/ena/data/view/PRJEB13687) using the Hisat2 short read aligner version 2.0.4 (Kim et al., 2015) and the Ensembl gene annotation (Labrus_bergylta.BallGen_V1.95, 11/25/2018, www.ensembl.org). Transcript abundance for the individual libraries was estimated using FeatureCounts (Liao et al., 2014) of the Subread package (http://subread.sourceforge.net/). Differential expression analysis was performed using the Bioconductor R package (version 3.4.4) DESeq2 (version 1.18.1) (Love et al., 2014). Genes of which fewer than five samples had gene counts below or equal to 10 reads were excluded from further analysis prior to normalization and differential expression analysis. Significantly expressed genes (p < 0.01) were used for further downstream analysis using the DAVID Bioinformatics Resources 6.8 (https://david.ncifcrf.gov/) with default settings (GO, UP_KEYWORDS and KEGG pathway analysis). Heat maps for hierarchical clustering of differentially expressed genes using multi/group comparison were embedded in the Qlucore omics explorer software package version 3.2 (Qlucore AB, Lund, Sweden).

## 3. Results

### 3.1. Properties of contraction types

The wrasse intestines reacted to the presence of a bolus of nutrients via dynamic changes in the intestinal diameter that reflected smooth muscle-driven contractions and relaxations of the gut. Depending on propagation distance and velocity, the contractile activity was classified as either standing contractions, ripples, or slow propagating contractions, according to the method described by Le et al. (2019a). Standing contractions are non-propulsive, while ripples and slow propagating contractions are two subtypes of propulsive contractions. A Kruskal-Wallis test on our data set showed that slow propagating contractions had higher amplitudes and lasted longer than other contraction types (slow propagating contractions > ripples > standing contractions, p < 0.0001) (Table S1). Ripples propagated for a shorter distance (p = 0.007) and at a higher velocity than slow propagating contractions (p < 0.0001).

### 3.2. Effect of nutrients on the time for emptying bulbous (BET)

Six types of bolus were inserted into the isolated intestines in order to analyze gut motility. However, since we were unable to block endogenous hydrolysis of intact protein and intact lipid, as previously mentioned we pooled data from intact and hydrolyzed lipid treatments into a group for lipid, and data from intact and hydrolyzed protein into a group for protein. The bulbous (Segment 1) emptying time (BET) was found to be 3.4±1.0 h in the lipid treatment and 3.9±1.5 h in the protein group (mean±s.d.) (Figure 2). In contrast, it only took between 0.6 and 2.5 h, with an average of 1.1±0.7 h, for a cellulose bolus to pass through Segment 1. The cellulose group had a faster BET than lipid (ANOVA, p = 0.003) and protein (ANOVA, p < 0.001) (Figure 2). The evacuation time for the plastic bead group was much more variable at between 0.6 and 5.9 hours, and was not different to the protein, lipid or cellulose groups.

**Figure 2.**
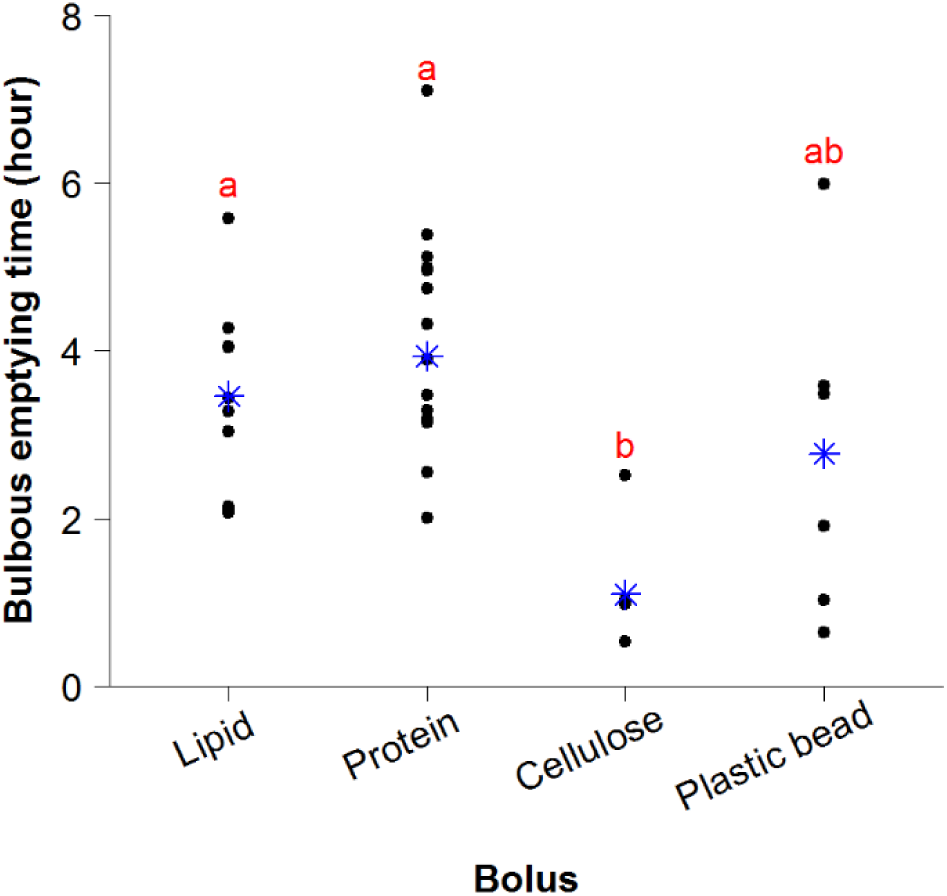
Evacuation time of Segment 1 (the bulbous). Analysis of intestines fed different nutrient boli (lipid, protein, cellulose or plastic bead) *in vitro*. Black dots refer to individual intestines, blue stars to mean for each treatment. Letters indicate significant differences in time to emptying of the bulbous between treatments (ANOVA followed by Tukey HSD; n = 11 for lipid, n = 12 for protein, n = 6 for cellulose or plastic bead).

### 3.3. Effects of diets on motility patterns

#### 3.3.1. Segment 1 (the bulbous)

Bolus composition had significant effects on frequency of contractions of all three motility patterns. When cellulose was present in Segment 1 (period I), it induced more standing contractions (1.5±0.5 contractions per min. for every mm gut length, cpm) compared to lipid (0.9±0.2 cpm) (lme, p = 0.03) and protein (0.64±0.05 cpm) (lme, p = 0.001). The frequency of standing contractions in the plastic bead group was 0.9±0.4 cpm and tended to be lower than that in the cellulose group (lme, p = 0.08). However, the rate of standing contractions in the plastic bead group was not different from that of the lipid and protein treatments (Figure 3 A). The presence of cellulose in Segment 1 also induced more propulsive contractions (0.26±0.11 cpm for slow propagating contractions and 0.08±0.02 cpm for ripples) compared with protein (0.09±0.04 for slow propagating contractions and 0.03±0.02 cpm for ripples) (lme, p < 0.05) (Figure 3 B and C). The frequency of slow propagating contractions in Segment 1 of intestines administered a lipid bolus did not differ from that in the protein group, but it was lower than in the cellulose group (lme, p = 0.04) (Figure 3 B). Ripples in the lipid group had the similar frequency to that in the cellulose group but compared to the protein treatment the contraction types occurred at a higher rate (lme, p = 0.04) (Figure 3 C). The frequency of propulsive contractions in the plastic bead treatments was not different from the three other groups (Figure 3 B and C). After ingesta left Segment 1 (period II), the frequency of all three contraction types declined in all four treatments (lme, p < 0.01) (black asterisk, Figure 3 A - C).

**Figure 3.**
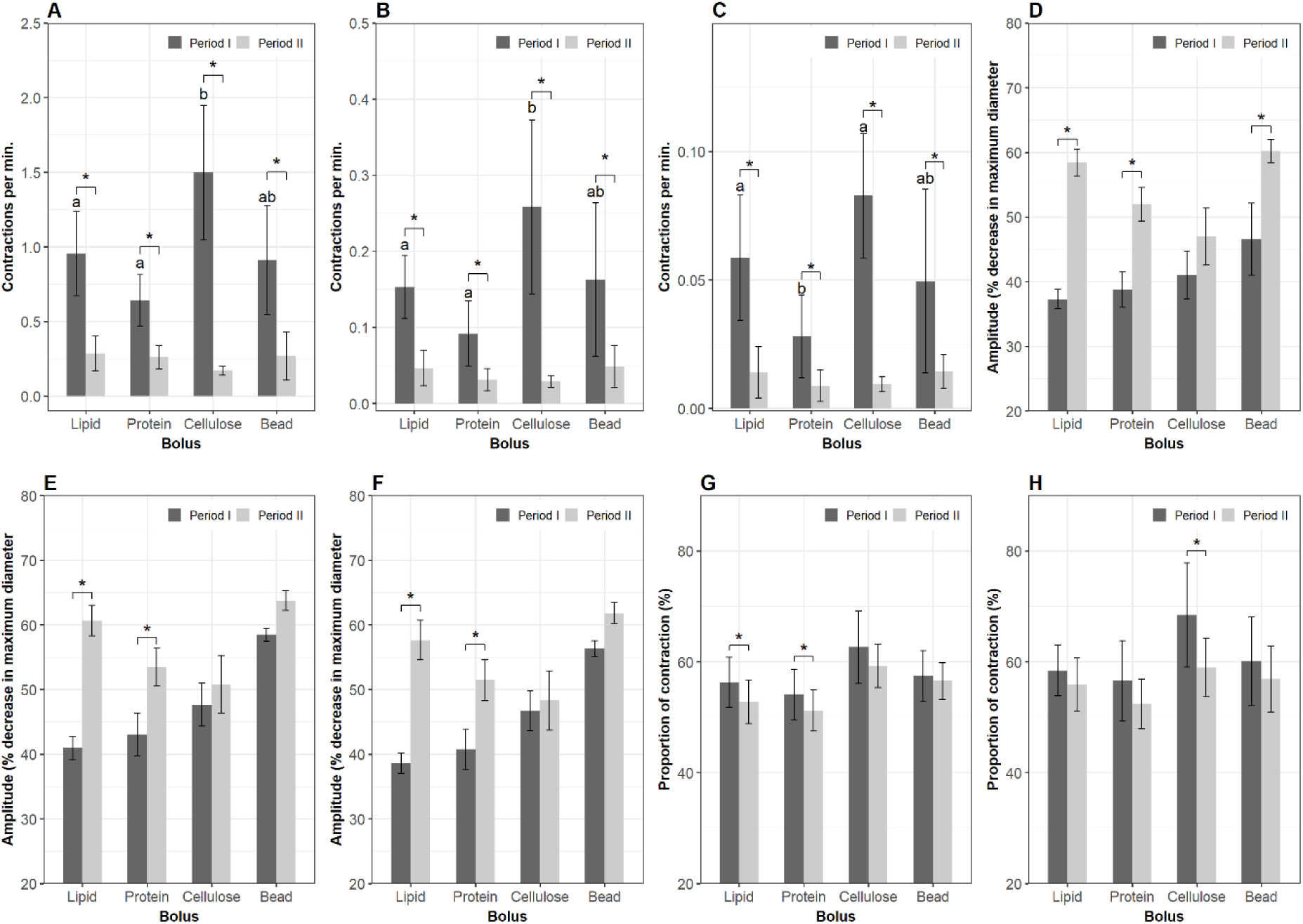
Motility parameters in Segment 1 (the bulbous). Contractions were analyzed in Segment 1 of the intestines when a nutrient bolus (lipid, protein, cellulose or plastic bead) was presented (period I) and after the bolus left (period II) this segment. Frequency (mean±s.d.) of standing contractions (A), slow propagating contractions (B) and ripples (C). Amplitude (median±s.e.m.) of standing contractions (D), slow propagating contractions (E) and ripples (F). Proportion (mean±s.d.) of slow propagating contractions (G) and ripples (H) which propagate to an antegrade direction. Significant differences (p < 0.05) between the four bolus treatments within period I are annotated by Latin letters and differences within period II by Greek letters. Asterisks and brackets show a significant difference (p < 0.05) between periods. Data were analyzed using linear mixed-effects models - lme test followed by Tukey HSD in R for (A - C); Generalized Linear Mixed Models via PQL - glmmPQL test for (D - H), with n = 11 for lipid, n = 12 for protein, n = 6 for cellulose or plastic bead.

The amplitude of all contraction types in Segment 1 was not different among the four bolus treatments during periods I and II. When ingesta exited the bulbous (period II) the amplitude increased in all contraction types in the lipid and protein group (lme, p < 0.001). However, this was not observed in the cellulose group. The plastic bead induced an increase in amplitude of standing contractions (lme, p = 0.0009) but no change in amplitude of propulsive contractions (Figure 3 D - F).

During the residence time of cellulose in Segment 1, 62.6±6.6% of the total slow propagating contractions and 68.4±9.4% of the total ripples propelled the bolus in an anterograde direction (direction towards the anus). The percentage of anterograde slow propagating contractions tended to be higher in the cellulose group than in the protein group (glmmPQL, p = 0.06) (Figure 3 G). Post-residence time in Segment 1, the proportion of anterograde slow propagating contractions was reduced in the lipid (glmmPQL, p = 0.007) and protein treatment groups (glmmPQL, p = 0.01). However, this parameter did not change in the cellulose or the plastic bead group (Figure 3 G). The percentage of anterograde ripples decreased after ingesta left the bulbous in the cellulose treatment (glmmPQL, p = 0.01) but did not change in the three other groups (Figure 3 H).

#### 3.3.2. Segment 2

The frequencies of the three contraction types in Segment 2 were not affected by bolus composition in either period I or II. However, the rate of contractions in Segment 2 was reduced when the boli had left Segment 1 (lme, p < 0.05) (Figure 4 A - C). Standing contractions and ripples had a higher amplitude in the plastic bead treatment compared to the lipid and the protein groups (glmmPQL, p < 0.05) during the residence time in Segment 1. The amplitude of contractions in the cellulose group did not differ from those of the three remaining treatments during either period I or II. During period II the amplitude of ripples in the lipid group was lower than in plastic bead group (glmmPQL, p = 0.04) (Figure 4 E). There was no change in amplitude of contractions in period II compared to that in period I regarding either bolus composition or contraction type. Propulsive contractions in an anterograde direction accounted for over 50% of the total propulsive contractions and the relative proportion of these contractions was not affected by nutrient treatments. The percentage of anterograde slow propagating contractions in Segment 2 decreased after plastic bead left Segment 1 (glmmPQL, p = 0.03) (Figure 4 C).

**Figure 4.**
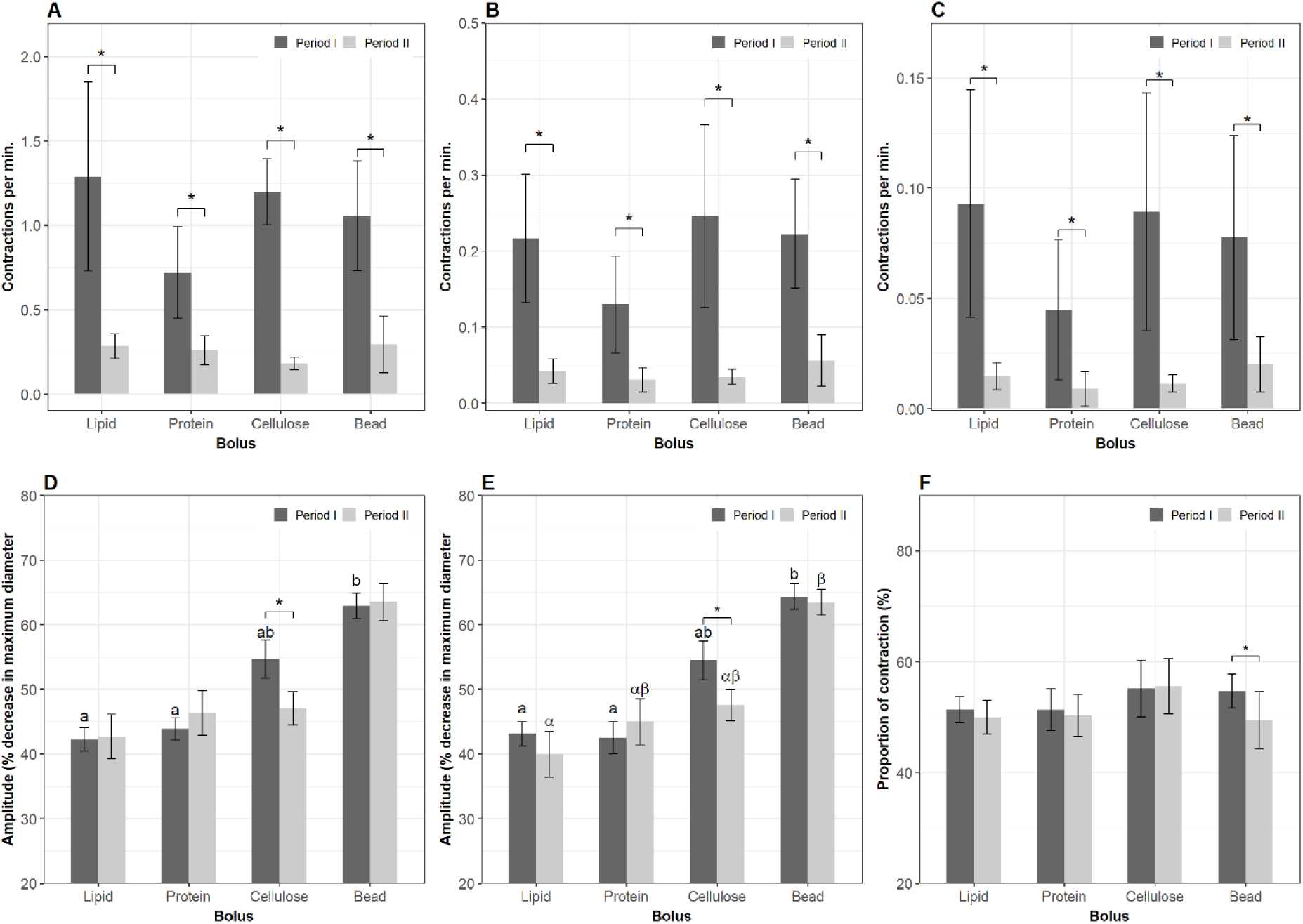
Motility parameters in Segment 2. Contractions were analyzed in Segment 2 of the intestines when a nutrient bolus (lipid, protein, cellulose or plastic bead) was presented (period I) and after the bolus left (period II) Segment 1. Frequency (mean±s.d.) of standing contractions (A), slow propagating contractions (B) and ripples (C). Amplitude (median±s.e.m.) of standing contractions (D) and ripples (E). Proportion (mean±s.d.) of slow propagating contractions (F) which propagate to an antegrade direction. Significant differences (p < 0.05) between the four bolus treatments within period I are annotated by Latin letters and differences within period II by Greek letters. Asterisks and brackets show a significant difference (p < 0.05) between periods. Data were analyzed using linear mixed-effects models - lme test followed by Tukey HSD in R for (A - C); Generalized Linear Mixed Models via PQL - glmmPQL test for (D - F), with n = 11 for lipid, n = 12 for protein, n = 6 for cellulose or plastic bead.

#### 3.3.3. Segment 3

Cellulose stimulated more standing contractions than protein in Segment 3 during period I (lme, p = 0.04) (Figure 5 A). Bolus composition did not affect the frequency of propulsive contraction types (Figure 5 B and C). The frequencies of the three contraction types diminished after ingesta left the Segment 1 in all four treatment groups (Figure 5 A - C). The evacuation of ingesta from Segment 1 produced a decrease in amplitude of all contraction types in the lipid and protein groups, and the amplitudes of contractions in these two groups were lower than in the plastic bead group. Amplitude of contraction in the cellulose group did not differ from the remaining groups either before or after ingesta left Segment 1 (Figure 5 D - F). The proportion of anterograde propulsive contractions did not differ between the four bolus groups in either period I or II.

**Figure 5.**
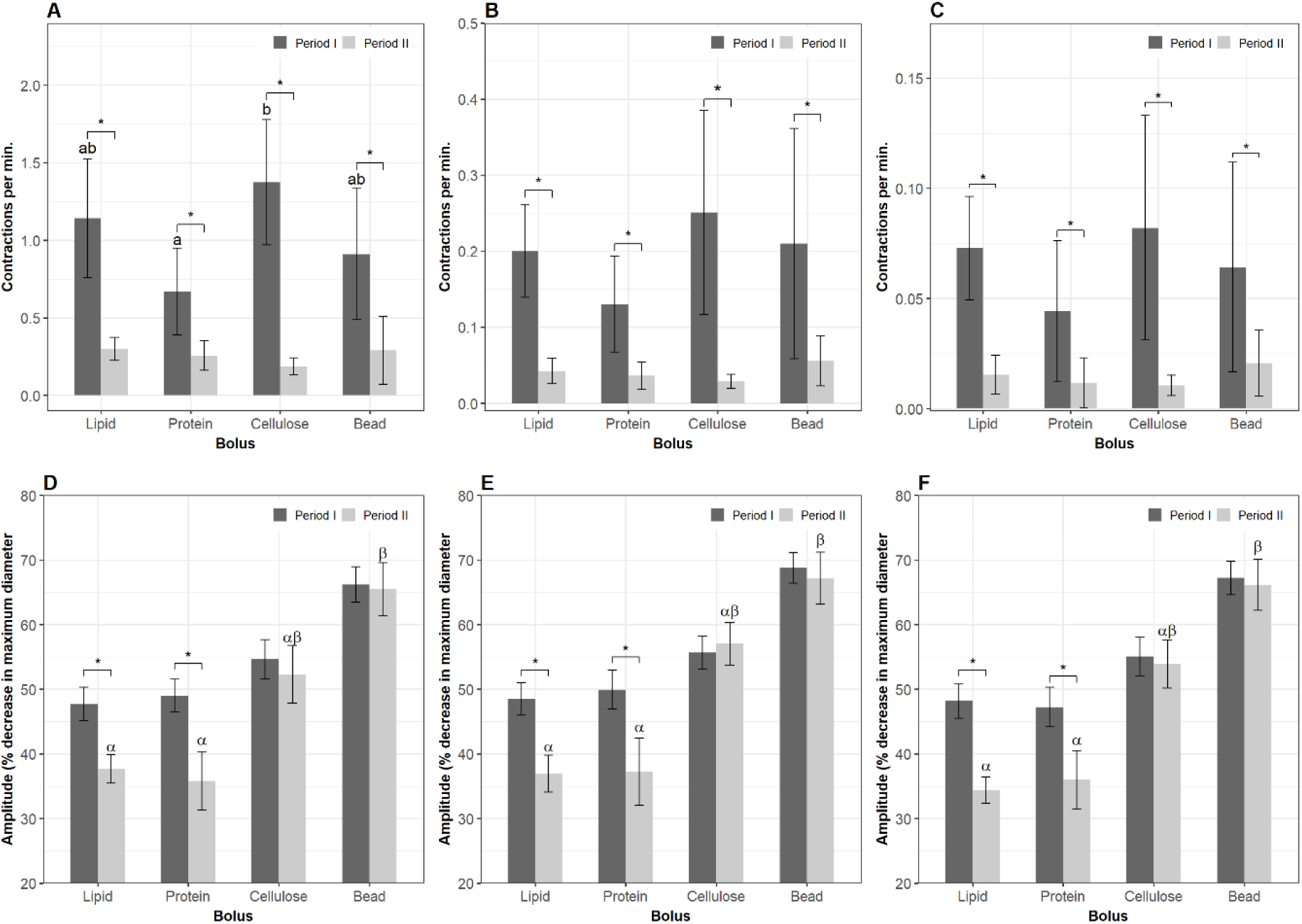
Motility parameters in Segment 3. Contractions were analyzed in Segment 3 of the intestines when a nutrient bolus (lipid, protein, cellulose or plastic bead) presented (period I) and after the bolus left (period II) Segment 1. Frequency (mean±s.d.) of standing contractions (A), slow propagating contractions (B) and ripples (C). Amplitude (median±s.e.m.) of standing contractions (D), slow propagating contractions (E) and ripples (F). Significant differences (p < 0.05) between the four bolus treatments within period I are annotated by Latin letters and differences within period II by Greek letters. Asterisks and brackets show a significant difference (p < 0.05) between periods. Data were analyzed using linear mixed-effects models - lme test followed by Tukey HSD in R for (A - C); Generalized Linear Mixed Models via PQL - glmmPQL test for (D - F), with n = 11 for lipid, n = 12 for protein, n = 6 for cellulose or plastic bead.

#### 3.3.4. Segment 4 (the hindgut)

During the residence time of the boli in Segment 1, no effects on the frequency of any of the three contraction types were observed in Segment 4. The evacuation of the boli out of Segment 1 suppressed the frequency of both non-propulsive and propulsive contractions in Segment 4 (Figure 6 A - C) (lme, p < 0.01). The rate of occurrence of slow propagating contractions in the cellulose group was lower than in the protein in period II (lme, p < 0.05) (Figure 6 B). The evacuation of ingesta out Segment 1 reduced the amplitude of standing contractions in three groups, but not in the plastic bead group (Figure 6 D); and amplitude of slow propagating contractions in the protein group (Figure 6 E); amplitude of ripples in the lipid and protein groups (Figure 6 F).

**Figure 6.**
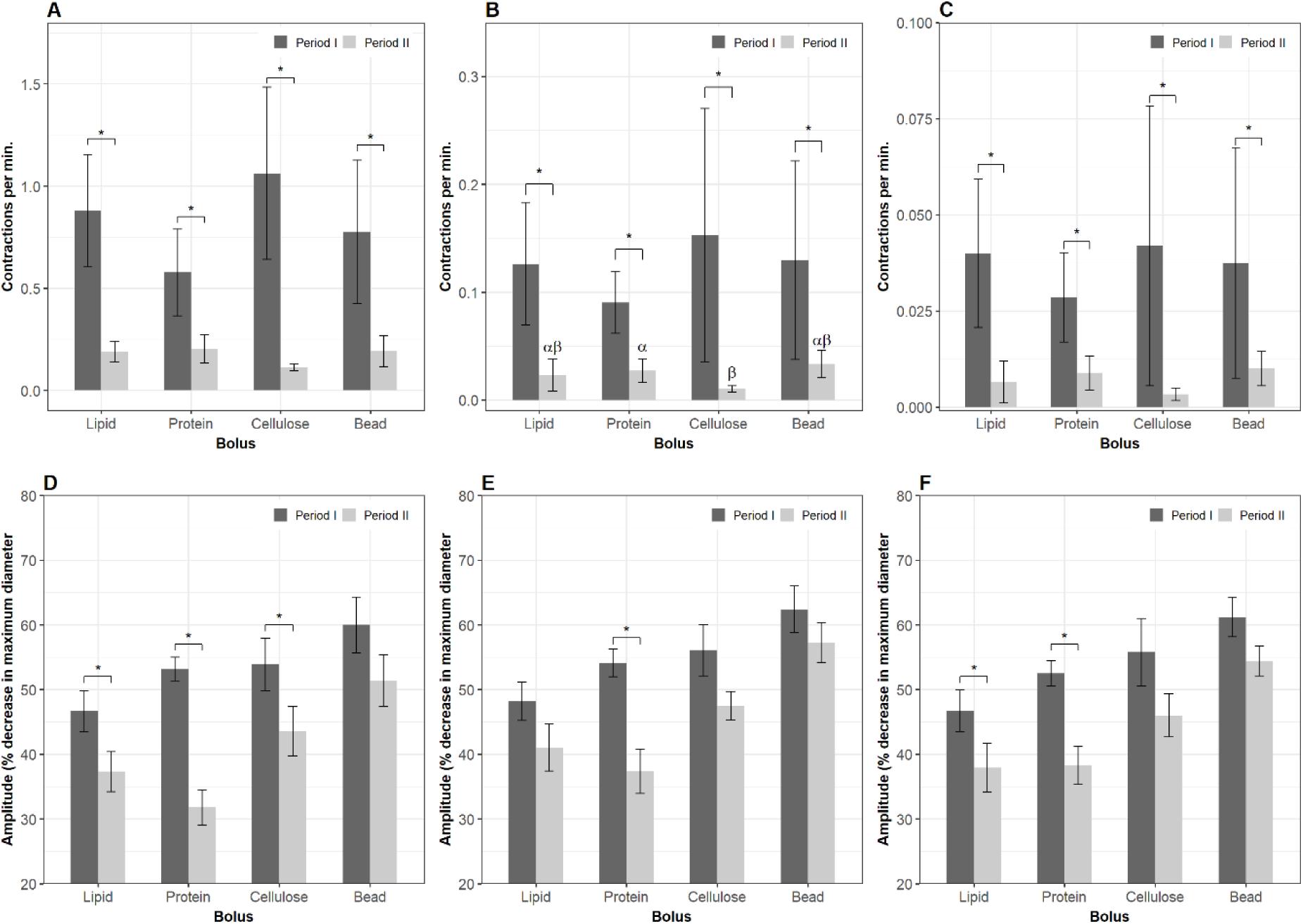
Motility parameters in Segment 4 (the hindgut). Contractions were analyzed in Segment 4 of the intestines when a nutrient bolus (lipid, protein, cellulose or plastic bead) presented (period I) and after the bolus leaves (period II) Segment 1. Frequency (mean±s.d.) of standing contractions (A), slow propagating contractions (B) and ripples (C). Amplitude (median±s.e.m.) of standing contractions (D), slow propagating contractions (E) and ripples (F). Significant differences (p < 0.05) between the four bolus treatments within period I are annotated by Latin letters and differences within period II by Greek letters. Asterisks and brackets show a significant difference (p < 0.05) between periods. Data were analyzed using linear mixed-effects models - lme test followed by Tukey HSD in R for (A - C); Generalized Linear Mixed Models via PQL - glmmPQL test for (D - F), with n = 11 for lipid, n = 12 for protein, n = 6 for cellulose or plastic bead.

#### 3.3.5. Comparing motility between the four intestinal segments

Duration and propagation velocity were not affected by bolus composition and did not change between period I and II. Variation in frequency of contractions and proportion of anterograde propagating contractions between the four intestinal segments were affected by bolus composition during period I (before ingesta left Segment 1). Overall, the frequencies of all contraction types in Segment 4 were lower than in the other three segments in all four treatment groups (Table 1).

**Table 1.** Frequency and direction of contractions between the four intestinal segments.

| Treatment | Contraction type | Frequency<sup>1</sup> | Anterograde direction<sup>2</sup> |
| --- | --- | --- | --- |
| Lipid | Standing contraction | $S1 = S4 < S2 = S3$ | - |
| | Slow propagating contraction | $S1 = S4 < S2 = S3$ | $S1 > S2 = S3 = S4$ |
| | Ripple | $S1 = S3 < S2$<br>$S2 > S3 > S4$<br>$S1 = S4$ | $S1 > S2 = S3 = S4$ |
| Protein | Standing contraction | $S1 = S2 = S3$<br>$S1 = S4$<br>$S2 = S3 > S4$ | - |
| | Slow propagating contraction | $S1 = S4 > S2 = S3$ | $S1 = S2 = S3 > S4$ |
| | Ripple | $S1 = S4 > S2 = S3$ | $S1 > S3 = S4$<br>$S1 = S2, S2 = S3$<br>$S2 > S4$ |
| Cellulose | Standing contraction | $S1 > S2 = S4$<br>$S1 = S3 > S4$<br>$S2 = S3$ | - |
| | Slow propagating contraction | $S1 = S2 = S3 > S4$ | $S1 > S2 = S3 > S4$ |
| | Ripple | $S1 = S2 = S3 > S4$ | $S1 = S2 = S3 = S4$ |
| Plastic bead | Standing contraction | $S1 = S2 = S3$<br>$S2 > S4$ | - |
| | Slow propagating contraction | $S1 = S4 < S2$<br>$S1 = S3, S2 = S3$ | $S1 = S2 = S3 = S4$ |
| | Ripple | $S1 = S4 < S2$<br>$S1 = S3, S2 = S3$ | $S1 = S2 = S3 = S4$ |

Variation in frequency of contraction and proportion of contractions propagating in an anterograde direction in the four intestinal segments before ingesta left Segment 1 (the bulbous).
<sup>1</sup> Frequency of contractions was analyzed using linear mixed models – lme.
<sup>2</sup> Proportion of contraction propagating in an anterograde direction to the total sum of contractions was analyzed using generalized linear mixed models (glmmPQL).
S1, S2, S3, and S4: Segment 1 (the bulbous), Segments 2, Segment 3 and Segment 4 (the hindgut).
=: No difference ( $p > 0.05$ ); < or >: less than or greater than ( $p \leq 0.05$ ).
N = 11 for lipid, 12 for protein, 6 for cellulose or plastic bead.

### 3.4. Effects of diets on gene expression

Gene expression in Segment 1 of the intestines fed cellulose was compared to that of empty intestines in order to elucidate the effect of stretch on gut activity. Effects of nutrient stimuli on gut metabolism were evaluated by analyzing differential gene expression in Segment 1 of the intestines between protein / lipid and cellulose groups.

#### 3.4.1. Differential gene expression

Genes were filtered using multi/group comparison (p < 0.01), which enables separation of the groups (protein, lipid, cellulose and empty) according to gene expression. Hierarchical clustering revealed two major clusters, one including the two “feed” treatment (protein and lipid) groups and the other one including empty intestines and cellulose-fed intestines. A total of 296 genes were clustered for this analysis (Figure 7). There were 178 genes that were differentially expressed (p < 0.01) in intestines fed with cellulose compared to empty intestines. A total of 628 and 275 were differentially expressed (p < 0.01) between protein versus cellulose, and lipid versus cellulose respectively (Table S2). Fewer genes were found to be differentially expressed when protein or lipid were administrated compared to the empty intestines.

**Figure 7.**
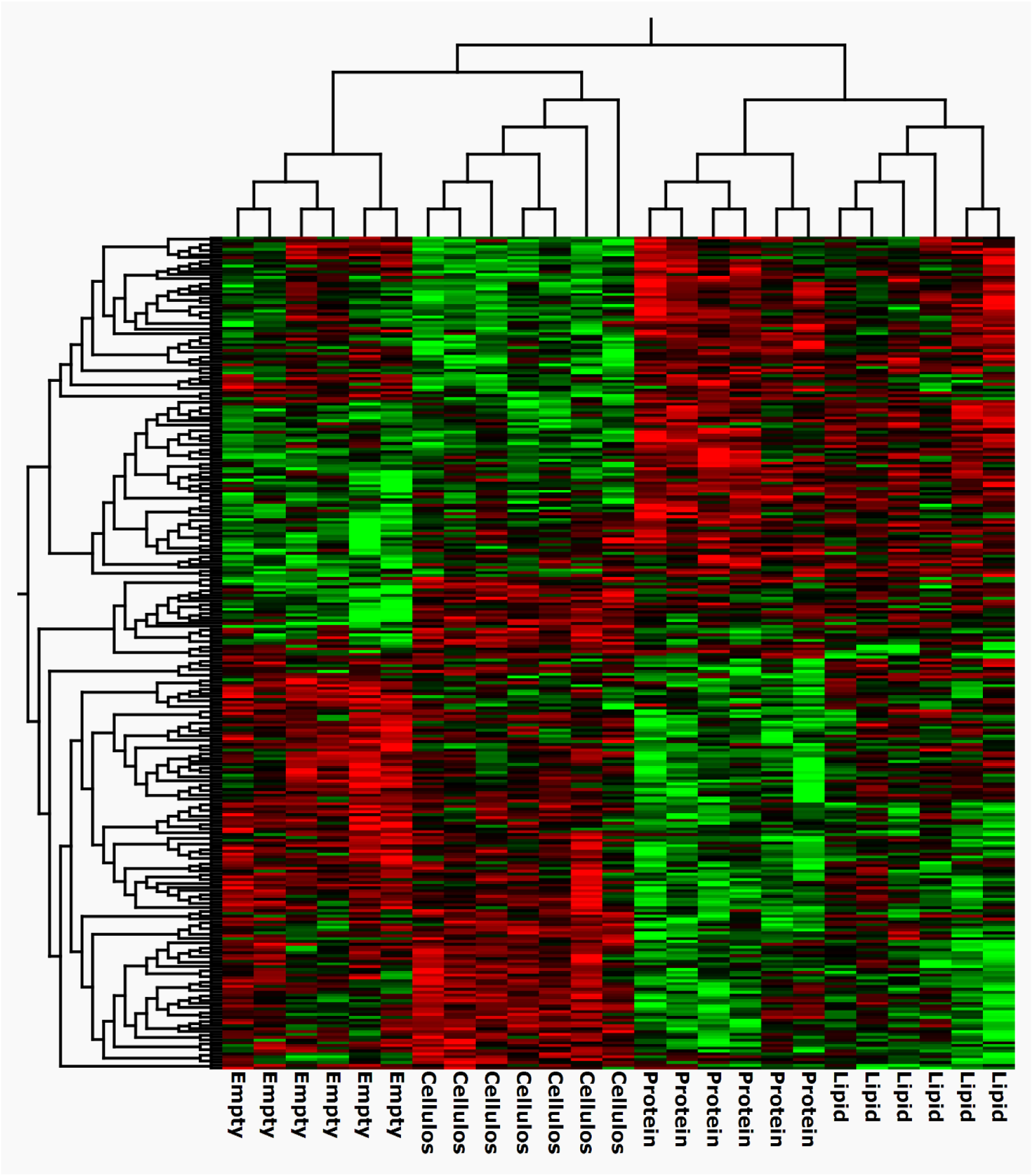
Nutrient sensing modulates the transcriptome in Segment 1. Hierarchical clustering of differentially expressed (p < 0.01) genes using multi/group comparison embedded in the Qlucore omics explorer. N = 6 for intestines fed with protein or lipid or empty intestines, n = 7 for intestines fed with cellulose.

Looking at overlapping genes in the DE gene lists, we found 135 genes to be unique for intestines administered cellulose and empty groups, while 43 genes were shared with protein and lipid treatments. The protein group shared 490 genes with the group, while lipid treatment had 153 genes in common with the cellulose treatment (Figure 8).

**Figure 8.**
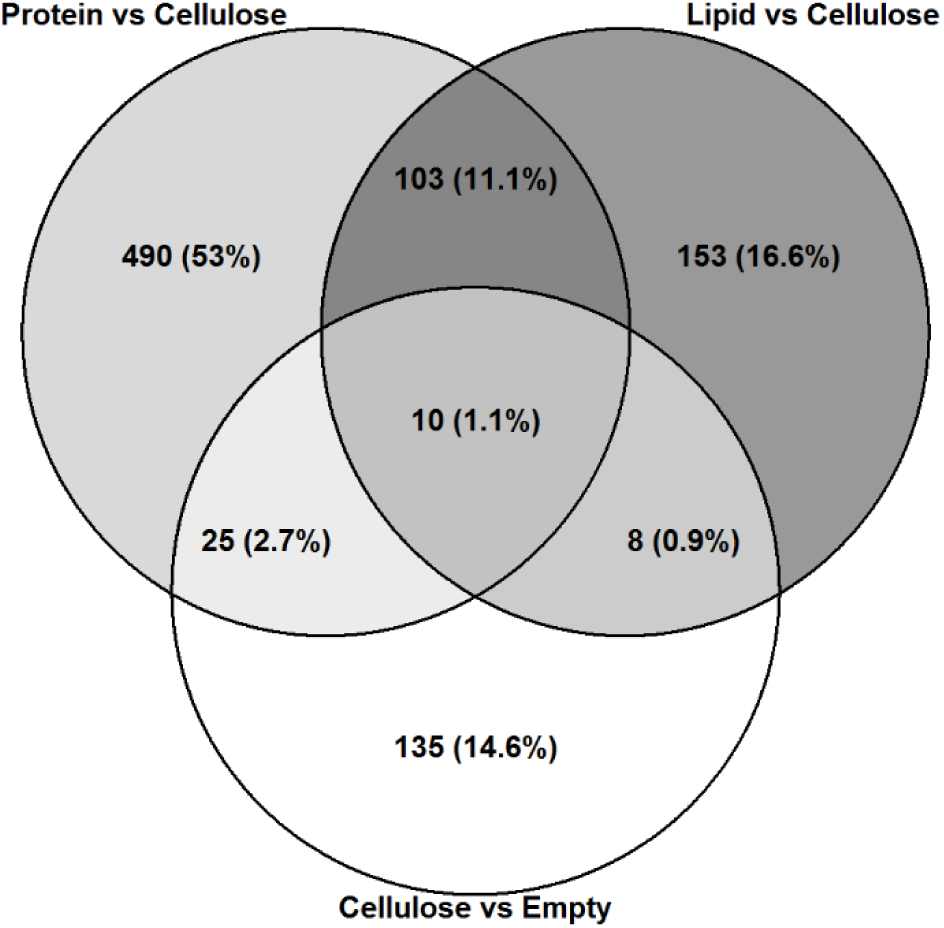
Nutrient sensing induces changes in gene expression in Segment 1. Venn diagram showing differentially expressed genes (p < 0.01) in intestines fed protein (n = 6) versus cellulose (n = 7), lipid (n = 6) versus cellulose, and cellulose versus empty groups following DESeq2 analysis of RNA seq data.

#### 3.4.2. Pathway and functional annotation analysis

Pathway analysis was performed using the differentially expressed genes of the different contrasts, including: cellulose versus empty, protein versus cellulose, and lipid versus cellulose. Enriched pathways related to calcium were observed in the cellulose versus empty comparison (FDR < 5%) but not in protein versus cellulose and lipid versus cellulose (Table 2).

**Table 2.** Pathways related to muscle contraction and nutrient sensing were regulated by diet bolus in Segment 1.

| Term | Count | % | p value | FDR (%) |
| --- | --- | --- | --- | --- |
| **Cellulose versus empty - upregulation** |  |  |  |  |
| Calcium | 13 | 17.1 | 6.4e-05 | 0.08 |
| GO:0005509~calcium ion binding | 12 | 15.8 | 1.4e-04 | 0.18 |
| **Protein versus cellulose - upregulation** |  |  |  |  |
| Secreted | 59 | 19.5 | 1.8e-07 | 2.5e-04 |
| GO:0008528~G-protein coupled peptide receptor activity | 5 | 1.7 | 2.0e-04 | 0.29 |
| GO:0007200~phospholipase C-activating G-protein coupled receptor signaling pathway | 7 | 2.3 | 8.5e-04 | 1.41 |
| Neuropeptide | 6 | 2.0 | 3.4e-05 | 0.05 |
| Opioid peptide | 3 | 1.0 | 6.4e-04 | 0.85 |
| GO:0005184~neuropeptide hormone activity | 5 | 1.7 | 1.54e-03 | 2.19 |
| GO:0001515~opioid peptide activity | 3 | 1.0 | 1.67e-03 | 2.36 |
| GO:0031628~opioid receptor binding | 3 | 1.0 | 1.67e-03 | 2.36 |
| GO:0006614~SRP-dependent cotranslational protein targeting to membrane | 41 | 13.5 | 1.8e-46 | 2.97e-43 |
| GO:0006364~rRNA processing | 53 | 17.5 | 1.0e-45 | 1.70e-42 |
| GO:0006413~translational initiation | 44 | 14.5 | 4.0e-43 | 6.69e-40 |
| GO:0019083~viral transcription | 41 | 13.5 | 9.3e-43 | 1.56e-39 |
| GO:0000184~nuclear-transcribed mRNA catabolic process, nonsense-mediated decay | 41 | 13.5 | 1.6e-41 | 2.72e-38 |
| Ribosomal protein | 43 | 14.2 | 6.3e-38 | 8.51e-35 |
| hsa03010:Ribosome | 43 | 14.2 | 8.0e-38 | 9.85e-35 |
| Ribonucleoprotein | 47 | 15.5 | 1.2e-33 | 1.61e-30 |
| GO:0003735~structural constituent of ribosome | 43 | 14.2 | 4.5e-32 | 6.40e-29 |
| GO:0005840~ribosome | 38 | 12.5 | 8.7e-32 | 1.13e-28 |
| GO:0022625~cytosolic large ribosomal subunit | 28 | 9.2 | 4.1e-31 | 5.33e-28 |
| GO:0006412~translation | 44 | 14.5 | 7.8e-31 | 1.31e-27 |
| GO:0044822~poly(A) RNA binding | 68 | 22.4 | 3.2e-20 | 4.65e-17 |
| GO:0022627~cytosolic small ribosomal subunit | 16 | 5.3 | 5.7e-16 | 7.22e-13 |
| GO:0003723~RNA binding | 37 | 12.2 | 2.7e-12 | 3.91e-09 |
| GO:0005925~focal adhesion | 23 | 7.6 | 4.0e-07 | 5.19e-04 |
| GO:0005829~cytosol | 88 | 29.0 | 9.3e-07 | 1.21e-03 |
| **Lipid versus cellulose - upregulation** |  |  |  |  |
| GO:0006364~rRNA processing | 10 | 9.2 | 4.8e-06 | 7.28e-03 |
| GO:0006413~translational initiation | 8 | 7.3 | 1.6e-05 | 2.49e-02 |
| GO:0006614~SRP-dependent cotranslational protein targeting to membrane | 7 | 6.4 | 1.9e-05 | 2.92e-02 |
| GO:0019083~viral transcription | 7 | 6.4 | 5.2e-05 | 0.08 |
| GO:0000184~nuclear-transcribed mRNA catabolic process,<br>nonsense-mediated decay | 7 | 6.4 | 7.3e-05 | 0.11 |
| Ribosomal protein | 7 | 6.4 | 4.4e-04 | 0.55 |
| hsa03010:Ribosome | 7 | 6.4 | 4.6e-04 | 0.55 |
| GO:0022625~cytosolic large ribosomal subunit | 5 | 4.6 | 5.7e-04 | 0.67 |
| GO:0006412~translation | 8 | 7.3 | 7.4e-04 | 1.12 |
| Ribonucleoprotein | 8 | 7.3 | 9.6e-04 | 1.20 |
| GO:0003735~structural constituent of ribosome | 7 | 6.4 | 1.86e-03 | 2.32 |
| GO:0005840~ribosome | 6 | 5.5 | 2.42e-03 | 2.79 |

The dashed borders separate clusters of the pathways which relate to the same physiological activity. FDR, False Discovery rate. N = 6 for intestines fed with protein or lipid or empty intestines, n = 7 for intestines fed with cellulose.

The largest number of significantly enriched pathways and functional annotations was observed for protein vs cellulose (FDR < 5%). Both the top enriched pathways (Table S3) and the largest cluster of pathways affected by protein (Table S4) were related to the level of ribosomal activity. The neuropeptide signaling pathway, opioid receptor activity pathways, inflammatory response, cytokine activity and G-protein coupled peptide receptor activity were among the many significantly enriched pathways upregulated by protein. Several of the pathways downregulated by protein were related to immune function, such as neutrophil chemotaxis, cellular response to interleukin-1, B cell receptor signaling pathway, and chemokine signaling pathway.

Many fewer significantly enriched pathways were observed in the lipid versus cellulose comparison. Similar to protein, genes involved in pathways related to ribosomal activity and RNA processing were enriched by lipid. Inflammatory response was also found to be significantly enriched by lipid (Table S4). Comparison of the expression of common genes in intestines between the protein and lipid groups found that inflammatory response, neuropeptide signaling pathway and ribosome were common pathways for the two treatments. The enriched pathways related to muscle contraction and nutrient sensing are shown in Table 2.

As identified by the pathway analysis, several genes coding for neuropeptide precursors were upregulated following protein and lipid treatment of the intestines (Table 3). None of these genes were differentially affected by cellulose compared to empty intestines. Other relevant neuropeptides such as proopiomelanocortin (*pomc*), neuropeptide Y (*npy*) and peptide YY (*pyy*) were filtered out before differential expression analysis, due to their low transcriptional levels.

**Table 3.** Genes coding for neuropeptides regulated by protein and lipid in Segment 1.

| Gene_ID | Gene_Name | Protein vs Cellulose |  | Lipid vs Cellulose |  |
| --- | --- | --- | --- | --- | --- |
|  |  | Fold change | Significance | Fold change | Significance |
| ENSLBEG00000013097 | *prepronociceptin (pnoca)* | 1.9 | * | 1.5 | - |
| ENSLBEG00000020351 | *neuromedin U (nmu)* | 2.5 | * | 1.3 | - |
| ENSLBEG00000019489 | *tachykinin 3 (tac3)* | 2.4 | ** | 1.5 | - |
| ENSLBEG00000015930 | *prodynorphin (pdyn)* | 1.6 | * | 1.5 | - |
| ENSLBEG00000009804 | *neuropeptide B (npb)* | 4.0 | *** | 2.2 | * |
| ENSLBEG00000014300 | *proenkephalin (penk)* | 1.8 | ** | 1.5 | * |
| ENSLBEG00000005406 | *cholecystokinin a receptor (cckar)* | 2.6 | * | 2.6 | * |
| ENSLBEG00000007237 | *cholecystokinin a (ccka)* | 1.3 | - | 1.3 | - |
| ENSLBEG00000024541 | *cholecystokinin b (cckb)* | -1.7 | - | -1.2 | - |

\* p<0.01, \*\*p<0.001, \*\*\*p<0.0001, - p ≥ 0.01
N = 6 for intestines fed with protein or lipid or empty intestines, n = 7 for intestines fed with cellulose.

## 4. Discussion

This study shows that boli of lipid and protein were evacuated from the bulbous (Segment 1) on an average of 3.4 to 3.9 h after they was administered to this section of ballan wrasse intestines *in vitro*. We have previously shown that more than 90% of the ingesta left the bulbous 4 h after the feed was administered *in vivo* (Le et al., 2019b). This suggest that an *in vitro* approach gives comparable results and may be used to estimate passage rate of ingesta in the alimentary tract.

The non-nutritive and indigestible cellulose was eliminated from the bulbous (Segment 1) more rapidly than protein and lipid. Slow propagating contractions may play a key role in propelling gut content since these may create a stronger force (Bharucha, 2012; Husebye, 1999; Kunze and Furness, 1999) and a longer propagation distance than other types of contraction (Brijs et al., 2014). Cellulose induced a higher frequency of slow propagating contractions than lipid and protein, resulting in a faster evacuation of the bolus out of the bulbous. Moreover, over 60% of the cellulose-induced slow propagating contractions were in an anterograde direction in the bulbous; and this value tended to be higher than that in protein and tended to be higher than in the lipid group. This suggests that the dominance of anterograde propulsive contractions aids in evacuating gut contents in the anal direction; hence, cellulose was propelled out of Segment 1 more rapidly than protein.

Cholecystokinin (CCK), a key hormone in the control of digestion, is released when food is present in the gut in both mammals (e.g., Grider, 1994; Liou et al., 2011; McLaughlin et al., 1998) and fish (e.g., MacDonald and Volkoff, 2009; Pitts and Volkoff, 2017). CCK performs its functions related to gastrointestinal motility and digestion mainly through Cholecystokinin A Receptor (CCKAR) in mammals [reviewed by Staljanssens et al. (2011)] and in fish (Tinoco et al., 2015; Zhang et al., 2017). We have previously found that in ballan wrasse, CCK works mainly through CCKA receptors to suppress propulsive contractions, thus prolonging the residence of food in the bulbous for optimal digestion (Le et al., 2019a). In the present study, we found that the presence of either lipid or protein in the lumen upregulated expression of *cckar* (CCKA receptor) which would be activated by CCK to suppress slow propagating contractions in Segment 1 to delay evacuation of the caloric ingesta from the bulbous. In contrast, there was no change in *cckar* expression in the bulbous of intestines administered cellulose compared to the empty intestines. Instead of activating CCKA receptor to slow the rate of evacuation, cellulose generates a different motility pattern, with increased frequency of propulsive contractions, predominantly anterograde slow propagating contractions that accelerate the elimination of the indigested content from Segment 1. The upregulation of *cckar* expression in Segment 1 by lipid and protein may be linked to the lower frequency of both ripples and slow propagating contractions in this segment compared to Segment 2. Conversely, there was no difference in the frequency of these contraction types between the three first segments in intestines fed with cellulose, which showed lower expression of *cckar* than those fed with lipid or protein.

Compared to cellulose, lipid and protein induced an increase in gene expression of *penk* and *npb* which suppress food intake (Aikawa et al., 2008; De Laet et al., 1989; Yang et al., 2014). This suggests that the presence of a caloric bolus in the gastrointestinal lumen stimulates enterocytes to secrete and send satiety signals to the central nervous system to inhibit feeding intake (Druce and Bloom, 2006; Rønnestad et al., 2017; Volkoff, 2016).

Lipid and protein had similar evacuation rates from the bulbous, and both were lower than cellulose. However, it appeared that the response of ballan wrasse intestines to the luminal presence of lipid somewhat differed from that of protein in the bulbous, with regard to both motility patterns and gene expression. Compared to protein, intestines fed lipid generated more ripples, which were suggested to have function in mixing rather than propelling gut content (D’Antona et al., 2001), to digest the hardly digestible fat. We have previously found that in ballan wrasse, only 50% of the lipid is digested in the bulbous and the digestive process needed to continue in the following sections (Le et al., 2019b), which resulted in the higher frequency of standing contractions to optimize the nutrient extraction in this segment (Gwynne and Bornstein, 2007b; Huizinga et al., 2014). Moreover, retrograde contractions increased in Segment 2 in the lipid group in order to propel the partly digested lipid bolus back to Segment 1, the main site of digestion and absorption in the ballan wrasse intestine (Le et al., 2019b). More than 70% of protein is digested and absorbed in the bulbous of ballan wrasse (Le et al., 2019b). It is therefore not necessary for Segments 2 and 3 to generate high frequencies of standing contractions for mixing and digesting (Schemann and Ehrlein, 1986) the already well-digested content received from the previous section. Also, intestines with a protein-bolus apparently do not need to increase anterograde contractions (i.e. the proportion of anterograde slow propagating contractions and ripples did not differ between the three first segments) to propel the well-digested protein back towards the bulbous.

Although they had similar evacuation rates, the difference in digestibility in the bulbous between lipid (50%) and protein (70%) (Le et al., 2019b) may correlate with the observed difference in regulation of some pathways between these two groups. Pathways related to regulation of opioid peptide and neuropeptides were enriched in the bulbous by protein but not by lipid. Prepronociceptin (PNOCA) and prodynorphin (PDYN) are prepropeptides which are proteolytically processed to form the various secreted opioid neurotransmitters including dynorphins, enkephalins, endorphins, endomorphins and nociceptin (Chavkin et al., 1985; LaForge et al., 2004). Most of these opiates, such as nociceptin, have been shown to inhibit gastrointestinal motility (Bueno and Fioramonti, 1988; Calo et al., 2000). The gut-brain peptide neuromedin U (NMU) has been reported to slow down gastric motility and emptying (Jarry et al., 2019). The upregulation of *pnoca, pdyn* and *nmu* in the bulbous of ballan wrasse intestines fed with protein (compared to cellulose) might be related to the lower frequency of all three contraction types in the protein than the cellulose group. Also, the expression of these three genes was altered by protein, but not by lipid (compared to cellulose), which might be related to the observed difference in motility patterns in the two first segments between the lipid and protein groups.

The presence of ingesta in the bulbous stimulate activity in the posterior segments, which resulted in the frequency of all contraction types in the four intestinal segments in period I being higher than that in period II (when the bulbous was empty). The inserted bolus in the bulbous was mixed and small amounts of gut content were propelled gradually to the next section until the bulbous was empty (Chang and Leung, 2014; Janssen et al., 2011; Welcome, 2018). Hence, role of the the high frequency of contractions in Segments 2 and 3 within period I was to continue the digestive process for the small amounts of ingesta which these segments received from the previous section. While the three first segments are mixing ingesta, Segment 4 – the hindgut had “cleaning” activity driven by the slow propagating contractions to eliminate undigested particles and waste (Der– Silaphet et al., 1998; Ehrlein et al., 1983) from previous meals and prepare for the remains of the current meal. When the bulbous was empty (period II), the frequency of all contraction types diminished in Segment 1 due to the absence of ingesta. In gastric vertebrates, the empty stomach will send hunger signals (Itoh et al., 1981; Szurszewski, 1969; Szurszewski, 1987). However, it is unknown how hunger is regulated in agastric vertebrates. The frequency of all contraction types also fell in the midgut (Segments 2 and 3) after Segment 1 was empty. This might be due to absorption of nutrients that did not require a high frequency of contractions and/or an artificial effect when energetic metabolites were gradually diminished after the long time since the intestines had been extracted from the fish body and incubated in medium (Cassim et al., 2017).

We used plastic beads to examine the stretch effect induced by an unbreakable and non-nutritive bolus on intestinal motility. The motility of the intestines containing the plastic beads could be classified into two categories: some intestines tended to propel the bead bolus to the distal intestine, while the others tended to reduce the anterograde/retrograde ratio of slow propagating contractions. The latter resulted in an extended residence time of beads in the bulbous and an increased amplitude of standing contractions, possibly in order to grind the beads into smaller particles (Schemann and Ehrlein, 1986). The presence of the hard bead in the bulbous may signal and drive Segments 2 and 3 to generate high-amplitude standing contractions that serve as a “physiological” brake (Chang and Leung, 2014) to delay the propulsion of the indigested meal to the distal sections. Overall, the amplitude of contractions in the plastic bead group was higher than that in other groups and this parameter did not differ between period I and II. This might be an artificial effect where an unbreakable bolus maintains its volume during its entire transit through the intestinal tract. Because of the stability in volume and shape which results in the reliable localization by video-scopy, beads have been widely used in studies of gastrointestinal transit and motility for several decades (e.g., Arkwright et al., 2016; Au - Hoffman et al., 2010; Camilleri and Linden, 2016; France et al., 2012; Koslo et al., 1986; Raffa et al., 1987; Reed et al., 2014). However, our results show that it is important to consider that the transit time and motility patterns induced by the beads differ significantly from the effect on these factors induced by a nutritious meal. Beads are therefore not a good proxy for intestinal motility during digestion.

Genes coding for calcium ion binding proteins were found to be enriched by administering either nutritive or non-nutritive boli. One of these genes, TRPM4 (Transient Receptor Potential Cation Channel Subfamily M Member 4), regulates smooth muscle contractions in a number of organs, including the intestine (Earley, 2013). This is probably related to the stretching effect induced by the presence of food. While the non-caloric cellulose bolus only induced mechanical factors, the caloric lipid and protein led to changes in a number of pathways related to cell activity. Most of the gastrointestinal hormones produced by enterocytes and other factors that regulate the digestion and absorption are peptides or proteins (Holmgren and Olsson, 2009; Lowe and Anderson, 2015; Welcome, 2018). Thus, protein and lipid boli upregulated a wide range of pathways related to the activity of ribosomes which, being protein factories (Green and Noller, 1997; Lengyel and Söll, 1969) in enterocytes might be to produce materials for secretion of enzymes, hormones, neurotransmitters and other components that are involved in the processes of digestion and absorption (Hall, 2013).

## 5. Conclusions

In conclusion, intestinal nutrient sensing regulates both motility patterns and gene expression to optimize digestion and absorption. The non-nutritive cellulose only triggered a stretching effect, increasing propulsive contractions to move the indigestible bolus from the bulbous to the hindgut. In contrast, the nutritive protein and lipid reduced the frequency of both non-propulsive and propulsive contractions to prolong the residence of the bolus in the bulbous, the main site of digestion in ballan wrasse. Protein and lipid upregulate *penk* and *npb*, which are known to send satiety signals to the central nervous system to suppress feeding. These nutrients also increased *cckar*, which is known to slow down the evacuation rate of the bulbous. Three other genes; *pnoca, pdyn* and *nmu*, were upregulated by protein but not by lipid. This may be related to the differences in motility patterns between the two nutrient groups.

## Author Contributions

Conceptualization, H.L., K.L., I.R., and Ø.S.; methodology, H.L., I.R., and Ø.S.; validation, H.L., K.L. and Ø.S.; formal analysis, H.L., K.L., A.E: and I.R.; investigation, H.L., K.L. A.E., and Ø.S.; resources, K.L.; data curation, K.L. and H.L.; writing—original draft preparation, H.L., K.L., and Ø.S.; writing—review and editing, H.L., K.L., I.R., and Ø.S.; visualization, H.L., K.L., I.R., and Ø.S.; supervision, Ø.S., I.R., and K.L.; project administration, Ø.S.; funding acquisition, Ø.S.

## Acknowledgments

We thank Skjalg Olsen and Jacob Wessels for their support in designing the *in vitro* organ system, Justine Giroud-Argoud for her help in dissecting fish, and Athanasios Sassalos for his support in protein gel and fatty acid analyses. We also thank Thomas E. Berentsen and Espen Grøtan (Marine Harvest) for supplying fish. The sequencing service was provided by the Norwegian Sequencing Centre (www.sequencing.uio.no), a national technology platform hosted by the University of Oslo and supported by the “Functional Genomics” and “Infrastructure” programs of the Research Council of Norway and the Southeastern Regional Health Authorities.

## Supplementary materials

**Table S1.** Parameters of three types of contractions

| Parameter | Contraction type | Min | Median | Max |
| --- | --- | --- | --- | --- |
| Amplitude (%) | Standing contraction | 7.7 | 47.4 | 77.9 |
|  | Ripples | 12.7 | 55.3 | 79.0 |
|  | Slow propagating contraction | 12.4 | 56.9 | 81.3 |
| Duration (s) | Standing contraction | 1.7 | 1.8 | 23.4 |
|  | Ripples | 1.79 | 7.2 | 258.2 |
|  | Slow propagating contraction | 1.72 | 15.3 | 506.6 |
| Distance (%) | Ripples | 1.2 | 2.2 | 16.6 |
|  | Slow propagating contraction | 1.3 | 2.5 | 16.1 |
| Velocity (mm s<sup>-1</sup>) | Ripples | 0.04 | 0.22 | 0.67 |
|  | Slow propagating contraction | 0.001 | 0.04 | 0.70 |

Data was collected in four segments of the intestines from a total of 35 individuals with n = 11 for lipid, n = 12 for protein, n = 6 for cellulose or plastic bead treatment in the Experiment 1.

**Table S2. Differential gene expression analysis between treatment groups**

**Table S3. Functional annotation enrichment analysis using DAVID**

**Table S4. Functional annotation clustering using DAVID**

**Figure S1.**
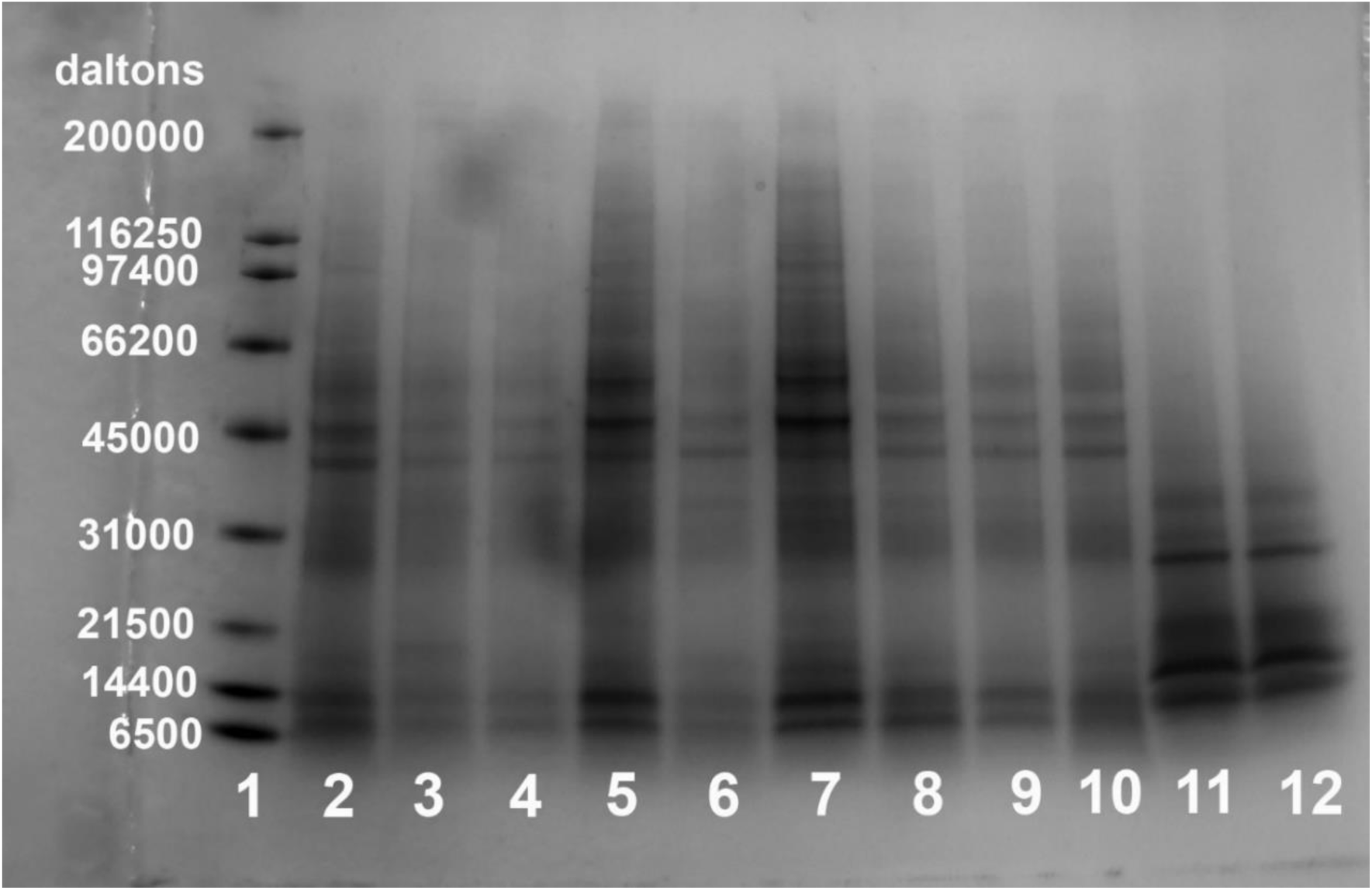
Protein gel electrophoresis of nutrient bolus and feces in Experiment 1. Lane 1, protein standard with multiple molecular weights (SDS-PAGE Molecular weight standards, Broad Range, Cat.No 161-0317, BIO-RAD); lanes 2, 4, 6, 8, and 10, feces collected from five individual intestines at 14 h after fed a bolus of intact protein; lanes 3, 5, 7, and 9, feces collected from four individual intestines at 14 h after fed a bolus of hydrolyzed protein; lanes 11 and 12, hydrolyzed protein bolus.

**Figure S2.**
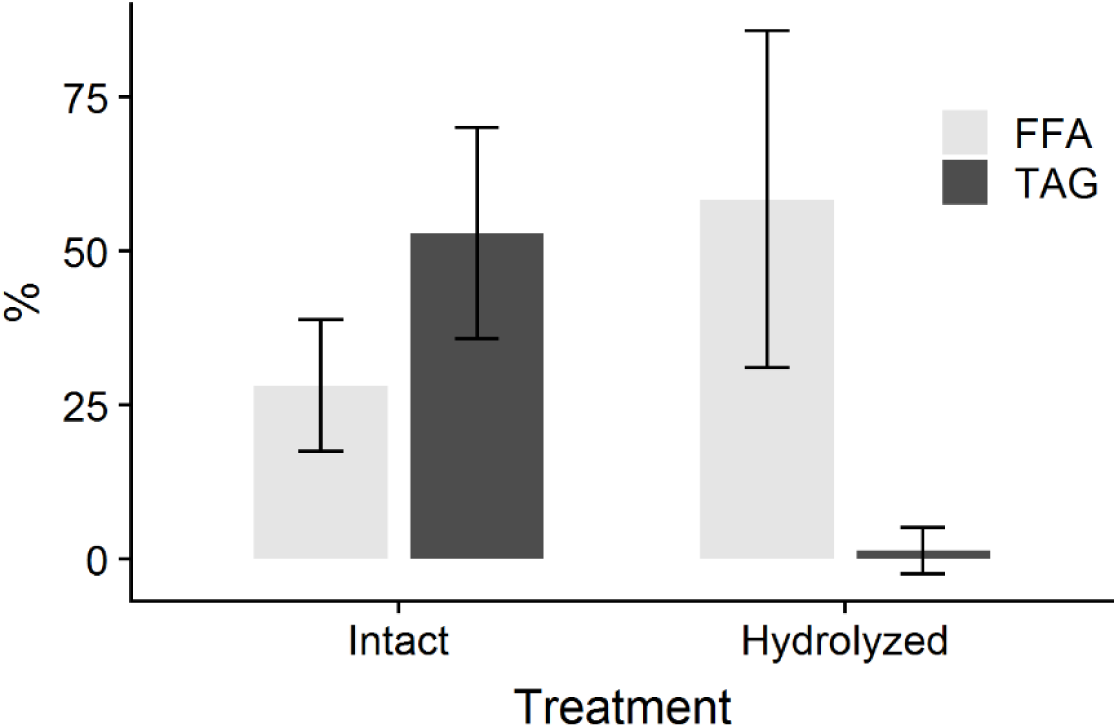
Lipid composition in feces collected from the isolated intestines at 14 h after the insert of a bolus in Experiment 1. FFA, free fatty acids; TAG, triacylglycerol.

## References

Food Forum; Food and Nutrition Board; Institute of Medicine. Relationships Among the Brain, the Digestive System, and Eating Behavior: Workshop Summary. Washington (DC): National Academies Press (US); 2015 Feb 27. 2, Interaction Between the Brain and the Digestive System. Available from: https://www.ncbi.nlm.nih.gov/books/NBK279994/.

Aikawa, S., Ishii, M., Yanagisawa, M., Sakakibara, Y. and Sakurai, T. (2008). Effect of neuropeptide B on feeding behavior is influenced by endogenous corticotropin-releasing factor activities. Regulatory Peptides 151, 147–152.

Andrews, P. L. R. and Young, J. Z. (1993). Gastric motility patterns for digestion and vomiting evoked by sympathetic nerve stimulation and 5-hydroxytryptamine in the dogfish *Scyliorhinus canicula*. Philosophical Transactions of the Royal Society of London, Series B: Biological Sciences 342, 363–380.

Arkwright, J. W., Underhill, I. D., Dodds, K. N., Brookes, S. J. H., Costa, M., Spencer, N. J. and Dinning, P. G. (2016). A composite fibre optic catheter for monitoring peristaltic transit of an intra-luminal bead. Journal of Biophotonics 9, 305–310.

Au - Hoffman, J. M., Au - Brooks, E. M. and Au - Mawe, G. M. (2010). Gastrointestinal Motility Monitor (GIMM). JoVE, e2435.

Babaei, S., Sáez, A., Caballero-Solares, A., Fernández, F., Baanante, I. V. and Metón, I. (2017). Effect of dietary macronutrients on the expression of cholecystokinin, leptin, ghrelin and neuropeptide Y in gilthead sea bream (*Sparus aurata*). General and Comparative Endocrinology 240, 121–128.

Bertrand, P. (2009). The cornucopia of intestinal chemosensory transduction. Frontiers in Neuroscience 3.

Bharucha, A. E. (2012). High amplitude propagated contractions. Neurogastroenterology and Motility 24, 977–982.

Borgstrom, S. and Arborelius, M., Jr. (1975). Influence of a fatty acid on duodenal motility. Scandinavian Journal of Gastroenterology 10, 599–601.

Brijs, J., Hennig, G. W., Axelsson, M. and Olsson, C. (2014). Effects of feeding on *in vivo* motility patterns in the proximal intestine of shorthorn sculpin (*Myoxocephalus scorpius*). The Journal of Experimental Biology 217, 3015–3027.

Brijs, J., Hennig, G. W., Gräns, A., Dekens, E., Axelsson, M. and Olsson, C. (2017). Exposure to seawater increases intestinal motility in euryhaline rainbow trout (*Oncorhynchus mykiss*). The Journal of Experimental Biology 220, 2397–2408.

Brijs, J., Hennig, G. W., Kellermann, A.-M., Axelsson, M. and Olsson, C. (2016). The presence and role of interstitial cells of Cajal in the proximal intestine of shorthorn sculpin (*Myoxocephalus scorpius*). The Journal of Experimental Biology 220, 347–357

Bueno, L. and Fioramonti, J. (1988). Action of opiates on gastrointestinal function. Bailliere’s Clinical Gastroenterology 2, 123–39.

Calo, G., Guerrini, R., Rizzi, A., Salvadori, S. and Regoli, D. (2000). Pharmacology of nociceptin and its receptor: a novel therapeutic target. British Journal of Pharmacology 129, 1261–1283.

Camilleri, M. and Linden, D. R. (2016). Measurement of gastrointestinal and colonic motor functions in humans and animals. Cellular and molecular gastroenterology and hepatology 2, 412–428.

Cannon, W. B. (1902). The movements of the intestines studied by means of the Röntgen rays. American Journal of Physiology-Legacy Content 6, 251–277.

Cannon, W. B. (1912). Peristalsis, segmentation, and the myenteric reflex. American Journal of Physiology-Legacy Content 30, 114–128.

Cassim, S., Raymond, V.-A., Lapierre, P. and Bilodeau, M. (2017). From in vivo to in vitro: Major metabolic alterations take place in hepatocytes during and following isolation. PLoS ONE 12, e0190366–e0190366.

Chang, E. B. and Leung, P. S. (2014). Gastrointestinal motility. In The Gastrointestinal System: Gastrointestinal, Nutritional and Hepatobiliary Physiology, (ed. P. S. Leung), pp. 35–62. Dordrecht: Springer Netherlands.

Chavkin, C., Shoemaker, W., McGinty, J., Bayon, A. and Bloom, F. (1985). Characterization of the prodynorphin and proenkephalin neuropeptide systems in rat hippocampus. The Journal of Neuroscience 5, 808–816.

Comesana, S., Velasco, C., Ceinos, R. M., Lopez-Patino, M. A., Miguez, J. M., Morais, S. and Soengas, J. L. (2018). Evidence for the presence in rainbow trout brain of amino acid-sensing systems involved in the control of food intake. American Journal of Physiology-Regulatory Integrative and Comparative Physiology 314, R201–R215.

Comesaña, S., Velasco, C., Conde-Sieira, M., Míguez, J. M., Soengas, J. L. and Morais, S. (2018). Feeding stimulation ability and central effects of intraperitoneal treatment of l-leucine, l-valine, and l-proline on amino acid sensing systems in rainbow trout: Implication in food intake control. Frontiers in Physiology 9.

Conde-Sieira, M. and Soengas, J. L. (2017). Nutrient sensing systems in fish: Impact on food intake regulation and energy homeostasis. Frontiers in Neuroscience 10, 603–603.

D’Antona, G., Hennig, G. W., Costa, M., Humphreys, C. M. and Brookes, S. J. H. (2001). Analysis of motor patterns in the isolated guinea-pig large intestine by spatio-temporal maps. Neurogastroenterology and Motility 13, 483–492.

De Laet, M.-H., Dassonville, M., Lotstra, F., Vierendeels, G., Rossier, J. and Vanderhaeghen, J.-J. (1989). Proenkephalin A associated peptides in the autonomic nervous system of the human infant gastrointestinal tract. Neurochemistry International 14, 129–134.

Defilippi, C. and Gómez, E. (1995). Effect of casein and casein hydrolysate on small bowel motility and D-Xylose absorption in dogs. Neurogastroenterology and Motility 7, 229–234.

Deloose, E., Janssen, P., Depoortere, I. and Tack, J. (2012). The migrating motor complex: control mechanisms and its role in health and disease. Nature Reviews Gastroenterology & Hepatology 9, 271–285.

Deloose, E. and Tack, J. F. (2015). Redefining the functional roles of the gastrointestinal migrating motor complex and motilin in small bacterial overgrowth and hunger signaling. American Journal of Physiology - Gastrointestinal and Liver Physiology.

Der–Silaphet, T., Malysz, J., Hagel, S., Arsenault, A. L. and Huizinga, J. D. (1998). Interstitial cells of Cajal direct normal propulsive contractile activity in the mouse small intestine. Gastroenterology 114, 724–736.

Druce, M. and Bloom, S. R. (2006). The regulation of appetite. Archives of Disease in Childhood 91, 183–187.

Earley, S. (2013). TRPM4 channels in smooth muscle function. Pflugers Archiv: European journal of physiology 465, 1223–1231.

Ehrlein, H. J., Reich, H. and Schwinger, M. (1983). Colonic motility and transit of digesta during hard and soft faeces formation in rabbits. The Journal of Physiology 338, 75–86.

Ehrlein, H. J., Schemann, M. and Siegle, M. L. (1987). Motor patterns of small intestine determined by closely spaced extraluminal transducers and videofluoroscopy. American Journal of Physiology - Gastrointestinal and Liver Physiology 253, G259–G267.

France, M., Bhattarai, Y., Galligan, J. J. and Xu, H. (2012). Impaired propulsive motility in the distal but not proximal colon of BK channel β1-subunit knockout mice. Neurogastroenterology and Motility 24, e450–e459.

Green, R. and Noller, H. F. (1997). Ribosomes and translation. Annual Review of Biochemistry 66, 679–716.

Grider, J. R. (1994). Role of cholecystokinin in the regulation of gastrointestinal motility. The Journal of Nutrition 124, 1334S–1339S.

Gwynne, R. M. and Bornstein, J. C. (2007a). Local inhibitory reflexes excited by mucosal application of nutrient amino acids in guinea pig jejunum. American Journal of Physiology - Gastrointestinal and Liver Physiology 292, G1660–G1670.

Gwynne, R. M. and Bornstein, J. C. (2007b). Mechanisms underlying nutrient-induced segmentation in isolated guinea pig small intestine. American Journal of Physiology - Gastrointestinal and Liver Physiology 292, G1162–G1172.

Gwynne, R. M., Thomas, E. A., Goh, S. M., Sjövall, H. and Bornstein, J. C. (2004). Segmentation induced by intraluminal fatty acid in isolated guinea-pig duodenum and jejunum. The Journal of Physiology 556, 557–569.

Hall, A. J. (2013). Chapter 57 - Small intestine. In Canine and Feline Gastroenterology, eds. R. J. Washabau and M. J. Day), pp. 651–728. Saint Louis: W.B. Saunders.

Hennig, G. W., Gregory, S., Brookes, S. J. H. and Costa, M. (2010). Non-peristaltic patterns of motor activity in the guinea-pig proximal colon. Neurogastroenterology and Motility 22, e207–e217.

Holmgren, S. and Olsson, C. (2009). Chapter 10 The neuronal and endocrine regulation of gut function. In Fish Physiology, vol. Volume 28 eds. N. J. Bernier G. V. D. Kraak A. P. Farrell and J. B. Colin), pp. 467–512. London: Academic Press.

Huge, A., Weber, E. and Ehrlein, H.-J. (1995). Effects of enteral feedback inhibition on motility, luminal flow, and absorption of nutrients in proximal gut of minipigs. Digestive Diseases and Sciences 40, 1024–1034.

Huizinga, J. D., Chen, J.-H., Zhu, Y. F., Pawelka, A., McGinn, R. J., Bardakjian, B. L., Parsons, S. P., Kunze, W. A., Wu, R. Y., Bercik, P. et al. (2014). The origin of segmentation motor activity in the intestine. Nature communications 5, 3326–3326.

Hurst, N. R., Kendig, D. M., Murthy, K. S. and Grider, J. R. (2014). The short chain fatty acids, butyrate and propionate, have differential effects on the motility of the guinea pig colon. Neurogastroenterology and Motility 26, 1586–1596.

Husebye. (1999). The patterns of small bowel motility: physiology and implications in organic disease and functional disorders. Neurogastroenterology and Motility 11, 141–161.

Itoh, Z., Aizawa, I. and Takeuchi, S. (1981). Neural regulation of interdigestive motor-activity in canine jejunum. American Journal of Physiology - Gastrointestinal and Liver Physiology 240, G324–G330.

Janssen, P., Vanden Berghe, P., Verschueren, S., Lehmann, A., Depoortere, I. and Tack, J. (2011). Review article: the role of gastric motility in the control of food intake. Alimentary Pharmacology & Therapeutics 33, 880–894.

Jarry, A. C., Merah, N., Cisse, F., Cayetanot, F., Fiamma, M. N., Willemetz, A., Gueddouri, D., Barka, B., Valet, P., Guilmeau, S. et al. (2019). Neuromedin U is a gut peptide that alters oral glucose tolerance by delaying gastric emptying via direct contraction of the pylorus and vagal-dependent mechanisms. FASEB Journal 33, 5377–5388.

Jiang, H. W., Bian, F. Y., Zhou, H. H., Wang, X., Wang, K. D., Mai, K. S. and He, G. (2017). Nutrient sensing and metabolic changes after methionine deprivation in primary muscle cells of turbot (*Scophthalmus maximus* L.). Journal of Nutritional Biochemistry 50, 74–82.

Keinke, O. and Ehrlein, H.-J. (1983). Effect of oleic acid on canine gastroduodenal motility, pyloric diameter and gastric emptying. Experimental Physiology 68, 675–686.

Kim, D., Langmead, B. and Salzberg, S. L. (2015). HISAT: a fast spliced aligner with low memory requirements. Nature Methods 12, 357.

Koslo, R. J., Burks, T. F. and Porreca, F. (1986). Centrally administered bombesin affects gastrointestinal transit and colonic bead expulsion through supraspinal mechanisms. Journal of Pharmacology and Experimental Therapeutics 238, 62–67.

Kunze, W. A. A. and Furness, J. B. (1999). The enteric nervous system and regulation of intestinal motility. Annual Review of Physiology 61, 117–142.

LaForge, K. S., Nyberg, F. and Kreek, M. J. (2004). Primary structure of guinea pig preprodynorphin and preproenkephalin mRNAs: multiple transcription initiation sites for preprodynorphin. Brain Research Bulletin 63, 119–126.

Le, H. T. M. D., Lie, K. K., Giroud-Argoud, J., Rønnestad, I. and Sæle, Ø. (2019a). Effects of cholecystokinin (CCK) on gut motility in the stomachless fish ballan wrasse (*Labrus bergylta*). Frontiers in Neuroscience 13.

Le, H. T. M. D., Shao, X., Krogdahl, Å., Kortner, T. M., Lein, I., Kousoulaki, K., Lie, K. K. and Sæle, Ø. (2019b). Intestinal function of the stomachless fish, ballan wrasse (*Labrus bergylta*). Frontiers in Marine Science 6, 1–15.

Lengyel, P. and Söll, D. (1969). Mechanism of protein biosynthesis. Bacteriological Reviews 33, 264–301.

Li, S., Li, J. B., Zhao, Y. L., Zhang, Q. and Wang, Q. C. (2017). Nutrient sensing signaling integrates nutrient metabolism and intestinal immunity in grass carp, *Ctenopharyngodon idellus* after prolonged starvation. Fish & Shellfish Immunology 71, 50–57.

Liao, Y., Smyth, G. K. and Shi, W. (2014). featureCounts: an efficient general purpose program for assigning sequence reads to genomic features. Bioinformatics 30, 923–930.

Liou, A. P., Lu, X., Sei, Y., Zhao, X., Pechhold, S., Carrero, R. J., Raybould, H. E. and Wank, S. (2011). The G-protein-coupled receptor GPR40 directly mediates long-chain fatty acid-induced secretion of cholecystokinin. Gastroenterology 140, 903-912.e4.

Love, M. I., Huber, W. and Anders, S. (2014). Moderated estimation of fold change and dispersion for RNA-seq data with DESeq2. Genome Biology 15, 550–550.

Lowe, J. S. and Anderson, P. G. (2015). Chapter 11 - Alimentary tract. In Stevens & Lowe’s Human Histology (Fourth Edition), eds. J. S. Lowe and P. G. Anderson), pp. 186–224. Philadelphia: Mosby.

MacDonald, E. and Volkoff, H. (2009). Neuropeptide Y (NPY), cocaine-and amphetamine-regulated transcript (CART) and cholecystokinin (CCK) in winter skate (*Raja ocellata*): cDNA cloning, tissue distribution and mRNA expression responses to fasting. General and Comparative Endocrinology 161, 252–261.

McLaughlin, J. T., Lomax, R. B., Hall, L., Dockray, G. J., Thompson, D. G. and Warhurst, G. (1998). Fatty acids stimulate cholecystokinin secretion via an acyl chain length-specific, Ca2+-dependent mechanism in the enteroendocrine cell line STC-1. The Journal of Physiology 513 (Pt 1), 11–18.

Miguel-Aliaga, I. (2012). Nerveless and gutsy: intestinal nutrient sensing from invertebrates to humans. Seminars in Cell & Developmental Biology 23, 614–620.

Olsson, C. and Holmgren, S. (2001). The control of gut motility. Comparative Biochemistry and Physiology Part A: Molecular and Integrative Physiology 128, 479–501.

Pitts, P. M. and Volkoff, H. (2017). Characterization of appetite-regulating factors in platyfish, *Xiphophorus maculatus* (Cyprinodontiformes Poeciliidae). Comparative Biochemistry and Physiology Part A: Molecular and Integrative Physiology 208, 80–88.

Raffa, R. B., Mathiasen, J. R. and Jacoby, H. I. (1987). Colonic bead expulsion time in normal and μ-opioid receptor deficient (CXBK) mice following central (ICV) administration of μ- and δ-opioid agonists. Life Sciences 41, 2229–2234.

Reed, D. E., Pigrau, M., Lu, J., Moayyedi, P., Collins, S. M. and Bercik, P. (2014). Bead study: a novel method to measure gastrointestinal transit in mice. Neurogastroenterology and Motility 26, 1663–1668.

Rønnestad, I., Gomes, A. S., Murashita, K., Angotzi, R., Jönsson, E. and Volkoff, H. (2017). Appetite-controlling endocrine systems in teleosts. Frontiers in Endocrinology 8, 73–73.

Rønnestad, I., Rojas-Garcia, C. R. and Skadal, J. (2000). Retrograde peristalsis; a possible mechanism for filling the pyloric caeca? Journal of Fish Biology 56, 216–218.

Sarna, S. K. (1986). Myoelectric correlates of colonic motor complexes and contractile activity. American Journal of Physiology - Gastrointestinal and Liver Physiology 250, G213–G220.

Schemann, M. and Ehrlein, H.-J. (1986). Postprandial patterns of canine jejunal motility and transit of luminal content. Gastroenterology 90, 991–1000.

Schmid, H.-R. and Ehrlein, H.-J. (1993). Effects of enteral infusion of hypertonic saline and nutrients on canine jejunal motor patterns. Digestive Diseases and Sciences 38, 1062–1072.

Schwartz, G. J. (2011). Gut fat sensing in the negative feedback control of energy balance — Recent advances. Physiology & Behavior 104, 621–623.

Siegle, M. L. and Ehrlein, H. J. (1988). Digestive motor patterns and transit of luminal contents in canine ileum. American Journal of Physiology - Gastrointestinal and Liver Physiology 254, G552–G559.

Spiller, R. C., Trotman, I. F., Adrian, T. E., Bloom, S. R., Misiewicz, J. J. and Silk, D. B. (1988). Further characterisation of the ‘ileal brake’ reflex in man-effect of ileal infusion of partial digests of fat, protein, and starch on jejunal motility and release of neurotensin, enteroglucagon, and peptide YY. Gut 29, 1042–1051.

Staljanssens, D., Azari, E. K., Christiaens, O., Beaufays, J., Lins, L., Van Camp, J. and Smagghe, G. (2011). The CCK(-like) receptor in the animal kingdom: Functions, evolution and structures. Peptides 32, 607–619.

Szurszewski, J. (1969). A migrating electric complex of canine small intestine. American Journal of Physiology-Legacy Content 217, 1757–1763.

Szurszewski, J. (1987). Physiology of the gastrointestinal tract. Electrical basis for gastrointestinal motility. New York: Raven, 383–422.

Tian, J., Wang, K. D., Wang, X., Wen, H., Zhou, H. H., Liu, C. D., Mai, K. S. and He, G. (2018). Soybean saponin modulates nutrient sensing pathways and metabolism in zebrafish. General and Comparative Endocrinology 257, 246–254.

Tinoco, A. B., Valenciano, A. I., Gómez-Boronat, M., Blanco, A. M., Nisembaum, L. G., De Pedro, N. and Delgado, M. J. (2015). Two cholecystokinin receptor subtypes are identified in goldfish, being the CCKAR involved in the regulation of intestinal motility. Comparative Biochemistry and Physiology Part A: Molecular and Integrative Physiology 187, 193–201.

van der Wielen, N., van Avesaat, M., de Wit, N. J. W., Vogels, J. T. W. E., Troost, F., Masclee, A., Koopmans, S.-J., van der Meulen, J., Boekschoten, M. V., Müller, M. et al. (2014). Cross-species comparison of genes related to nutrient sensing mechanisms expressed along the intestine. PLoS ONE 9, e107531–e107531.

Volkoff, H. (2016). The neuroendocrine regulation of food intake in fish: A review of current knowledge. Frontiers in Neuroscience 10.

Welcome, M. O. (2018). Gastrointestinal motor function. In Gastrointestinal physiology: Development, principles and mechanisms of regulation, pp. 353–453. Cham: Springer International Publishing.

Xu, D., He, G., Mai, K., Zhou, H., Xu, W. and Song, F. (2016). Postprandial nutrient-sensing and metabolic responses after partial dietary fishmeal replacement by soyabean meal in turbot (*Scophthalmus maximus* L.). British Journal of Nutrition 115, 379–388.

Yang, L., Sun, C. and Li, W. (2014). Neuropeptide B in Nile tilapia *Oreochromis niloticus*: Molecular cloning and its effects on the regulation of food intake and mRNA expression of growth hormone and prolactin. General and Comparative Endocrinology 200, 27–34.

Zhang, X., Tang, N., Qi, J., Wang, S., Hao, J., Wu, Y., Chen, H., Tian, Z., Wang, B., Chen, D. et al. (2017). CCK reduces the food intake mainly through CCK1R in Siberian sturgeon (*Acipenser baerii* Brandt). Scientific Reports 7, 12413.

